# AAV delivery of RNA editing machinery rescues SUDEP and seizure phenotype in a mouse model of Dravet Syndrome

**DOI:** 10.1101/2025.07.18.664895

**Authors:** Ellie M. Chilcott, Juan F. Antinao Diaz, Amanda Almacellas Barbanoj, Marc Moore, Anna Keegan, Zakary Waddington, Vicky Fang, Stephanie Schorge, Gabriele Lignani, Simon N Waddington, Rajvinder Karda

## Abstract

Dravet syndrome (DS) is a severe childhood genetic epilepsy, caused by *de novo* heterozygous mutations in *SCN1A*, resulting in a loss-of-function of the voltage-gated sodium ion channel, Nav1.1. Nav1.1 is expressed in the brain, and at a lower level, in the heart. DS manifests in the first year of life. Patients exhibit tonic-clonic seizures, febrile seizures, cognitive decline, developmental delays, ataxia, and sudden unexpected death from epilepsy (SUDEP). We have developed a novel AAV-F mediated CRISPR-Cas-inspired RNA targeting system (CIRTS) preclinical treatment to increase endogenous *Scn1a* and ameliorate the disease phenotype in a clinically-relevant heterozygous loss-of-function mouse model of DS. We designed novel guide RNAs (gRNAs) to target the long non-coding RNA, (or natural antisense transcript) of *Scn1a* to increase the expression of *Scn1a* mRNA in DS mice. We show that intracerebroventricular and intravenous administration of AAV-F-CIRTS-gRNA9 to target the brain and the heart to neonatal *Scn1a^+/-^*mice resulted in a significant increase in survival, and a reduction in SUDEP, febrile seizures and seizure duration. These findings provide proof-of-concept evidence that an AAV-F-CIRTS mediated therapy hold promise as a potential treatment for DS.

## INTRODUCTION

Dravet syndrome (DS) is a severe childhood genetic epilepsy, with an incidence rate of 1 in 15,400-40,900 live births worldwide(*1, 2*). The disease typically manifests within the first year of life accompanied by tonic-clonic seizures, febrile and afebrile seizures, episodes of status epilepticus, cognitive impairment, ataxia and unfortunately, ∼15% of patients experience sudden unexpected death in epilepsy (SUDEP)(*3–5*). >90% of patients harbour *de novo* loss-of-function variants in the *SCN1A* gene which encodes an α-subunit of a voltage-gated ion channel, Nav1.1(*6*). In humans, Nav1.1 expression has been detected in the hippocampus (including the dentate gyrus, CA3, and CA2 regions), layers V and VI of the cortex, the granular and molecular layers, as well as in the deep nuclei and Purkinje cells of the cerebellum(*7*) and the heart. Previous studies have explored altered cardiac function in individuals with DS and in mouse models, suggesting a potential contribution to SUDEP(*8–11*).

Current anti-seizure medications, including valproate, stiripentol, fenfluramine and cannabidiol can reduce the seizure frequency(*12*), but the disease manifestation continues. Thus, there remains a significant unmet need to establish an effective long-term treatment for DS.

Adeno-associated viral (AAV) vectors, specifically AAV9, have been used extensively for many preclinical(*13, 14*) and clinical trials(*15, 16*). However, the high doses of AAV9 required to achieve efficacious peripheral delivery has also caused liver toxicity and thrombotic microangiopathy(*16, 17*). As such, there is a need to develop novel capsids that enable efficient therapeutic delivery at a lower dose, avoiding immunotoxicity associated with the gene therapy vector. Novel AAV capsids targeting the central nervous system (CNS) have been generated(*18, 19*). One example of this is AAV-F, a novel capsid derived from AAV9, which has shown a greater CNS transduction efficiency relative to the parental AAV9 capsid(*18*) and has been used successfully in a preclinical study to treat a severe mitochondrial disorder(*20*).

RNA targeting technologies have emerged as therapeutic, inspired by successes in oligonucleotide based therapies(*21, 22*). Targeting RNA is advantageous over DNA as it can influence gene expression without permanent alterations to the host genome and modulate protein expression regardless of the specific mutation. Furthermore, DNA and RNA targeting systems such as Cas9 and Cas13 are built from bacterial proteins and have shown to induce an immune response *in vivo*(*23*). A new RNA editing technology has been developed to target RNA sequences, referred to as CRISPR-Cas-inspired RNA targeting system (CIRTS)(*24*). CIRTS is built from human proteins and previous validation studies *in vitro* and *in vivo* have shown the CIRTS system to modulate protein expression by RNA editing (*24–26*), although until now this has not been used in a therapeutic setting. In this study, we used a CIRTS construct consisting of a ssRNA binding protein (β-defensin 3), RNA hairpin-binding protein (TBP6.7) and effector protein (YTHDF2)(*24*) to down-regulate target RNA expression.

Here, we used the AAV-F-mediated CIRTS technology and designed novel guide RNAs (gRNAs) to edit a *Scn1a* specific long non-coding RNA (lncRNA) of the natural antisense transcript (NAT), which normally functions to negatively regulate *Scn1a* expression(*27*). In doing so, we aimed to ‘inhibit the inhibitor’, leading to an upregulation of *Scn1a* mRNA (Fig. 1A). We screened 21 gRNAs *in vitro* and selected two lead candidate gRNAs (gRNA9 & 21) which significantly increased endogenous *Scn1a* mRNA and shared highest homology to the human lncRNA sequence. We used a clinically-relevant DS heterozygous (*Scn1a*^+/-^) model, which recapitulates much of the human disease, including spontaneous and febrile seizures, and SUDEP(*28, 29*). We administered our candidates AAV-F-CIRTS-gRNA 9 & 21 vectors to neonatal DS mice via intracerebroventricular (ICV) and intravenous (IV) administration to target the brain and the heart. Candidate AAV-F-CIRTS-gRNA9 resulted in significant increase in survival over 100 days post-injection. In addition, febrile and spontaneous seizure assessment of DS mice injected with AAV-F-CIRTS-gRNA9 showed a significant reduction in seizure severity and seizure duration. Molecular analysis revealed a significant up-regulation of endogenous *Scn1a* mRNA in treated AAV-F-CIRTS-gRNA9 DS mice compared to untreated DS control mice. Our work demonstrates for the first time, a potential AAV-F mediated RNA editing approach to treat patients with DS.

**Fig. 1.**
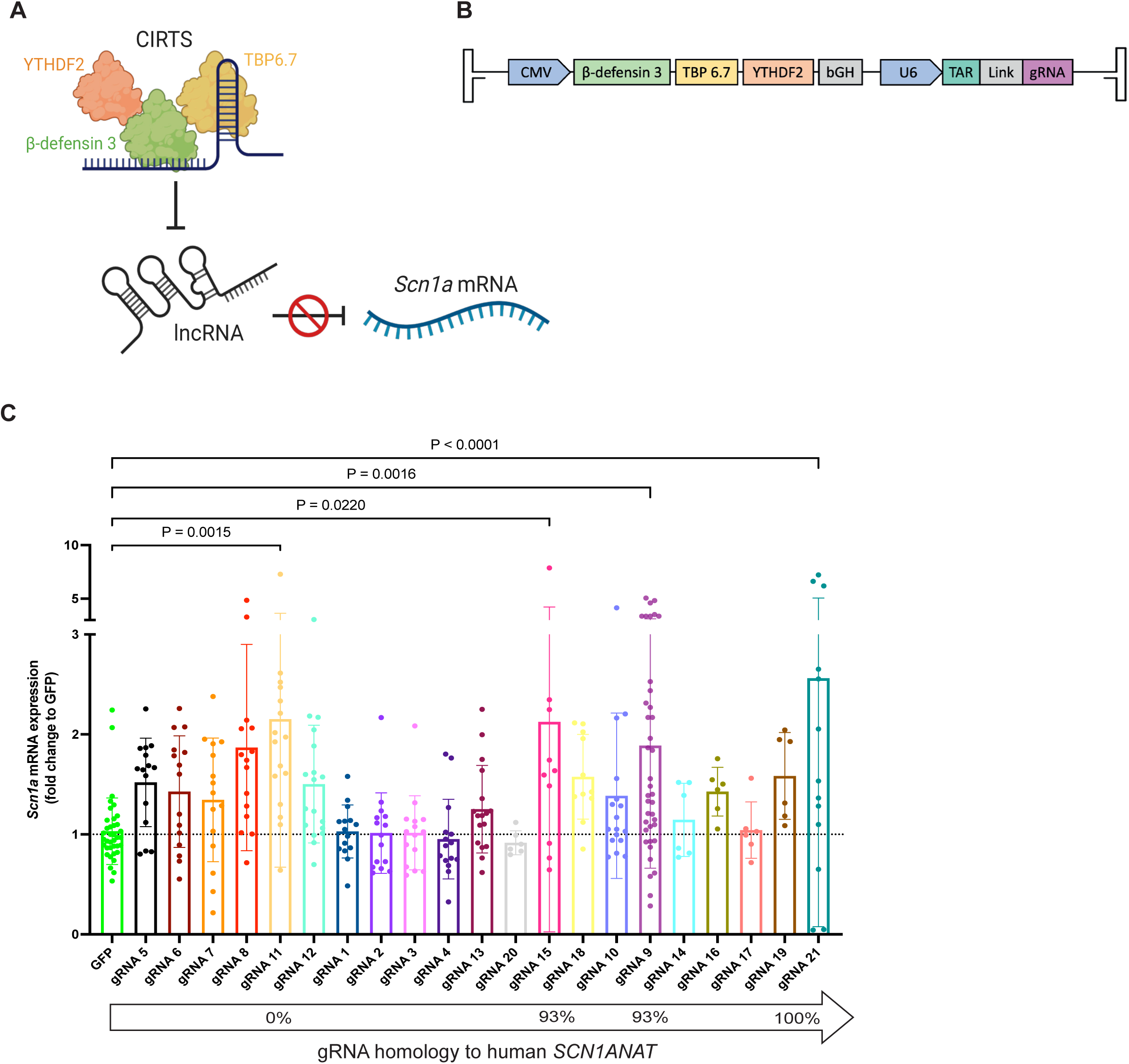
Novel gRNAs increase in endogenous *Scn1a in vitro*. (A) Schematic illustrating mechanism of action of CIRTS to increase endogenous *Scn1a* expression through targeting of the lncRNA. (B) Schematic diagram of the AAV-CIRTS-gRNA construct. (C) 4 novel AAV-CIRTS-gRNA constructs significantly increased endogenous *Scn1a* mRNA after transient transfection of differentiated mouse Neuro2a cells. The homology of these gRNAs to the human *SCN1ANAT* sequence are indicated on the graph as a %, with gRNAs ranked from least to most homologous. One-way ANOVA with Dunnett’s multiple comparisons.

## RESULTS

### Design and *in vitro* validation of novel gRNAs targeting *Scn1a* lncRNA

We firstly validated the CIRTS machinery and consistent with previous studies(*24*), AAV-CIRTS-gRNAluc was able to significantly reduce luciferase expression (Fig. S1). We designed 21 novel gRNA sequences (20-30mer) to target the *Scn1a* lncRNA (Fig. 1A) (here referred to as *Scn1anat*), which were cloned into the AAV-CIRTS plasmid (Fig. 1B). We validated our gRNA designs *in vitro*, in differentiated mouse Neuro2a cells and assessed the effects on endogenous *Scn1a* mRNA expression. Transient transfection of AAV-CIRTS-gRNA constructs onto differentiated Neuro2a cells resulted in a significant increase of endogenous *Scn1a* mRNA by 4 out of 21 AAV-CIRTS-gRNA constructs (Fig. 1C). Constructs containing gRNAs 9, 11, 15 and 21 led to average *Scn1a* mRNA fold changes of 1.89 (P=0.0016), 2.15 (P=0.0015), 2.12 (P=0.022) and 2.56 (P<0.0001), respectively. gRNAs 9 and 21 were selected for the *in vivo* study as they showed the highest homology to the *SCN1A* lncRNA (*SCN1ANAT*; 93.3% and 100%, respectively) (Fig. 1C).

### AAV-F-mediated RNA editing in DS mice significantly improves survival

In this study we crossed the males from 129Sv-*Scn1a^tm1Kea^*/Mmjax model(*28, 29*) with wild-type CD1 females to generate F2 129Sv-*Scn1a^tm1Kea^*/Mmjax x CD1 heterozygote females, which were then bred to wild-type C57BL/6J males to obtain experimental litters (*Scn1a*^+/-^). This enabled a larger litter size. *Scn1a*^+/-^ mice exhibit spontaneous seizures and SUDEP with an overall survival of 38.2% over 100 days of development (Fig. S2A), febrile seizures (Fig. S2D) and a reduction in *Scn1a* gene expression, particularly in the cortex at P50 (P=0.0078; Fig. S2E), consistent with previous studies(*22, 28, 30*). RNA-seq of *Scn1a*^+/+^ and *Scn1a^+/-^* cortical tissue at P50 confirmed 140 up- and 41 down-regulated genes (Fig. S3A and B), with significant down-regulation of *Scn1a* (−0.77Log2FC, P=5.85×10^-6^; Fig. S2C). Molecular function gene ontology (GO) analysis of the differentially expressed genes in *Scn1a^+/-^* mice reveal up-regulation in terms related to neuropeptide receptor activity, monoatomic ion transport and voltage-gated sodium channel activity (Fig. S3D).

Using this DS mouse model, we sought to determine the optimal capsid and gRNA combination by generating AAV-F and AAV9 vectors for AAV-CIRTS-gRNA9 and AAV-CIRTS-gRNA21, using AAV9 as a gold-standard comparison. At P0, *Scn1a*^+/-^ mice received an intracerebroventricular (ICV) and intravenous (IV) delivery of either AAV-F (7×10^10^ vg/pup) or AAV9 (7×10^11^ vg/pup), each encoding either gRNA9 or 21 to determine the effects on survival. This study was fully blinded and randomised (Fig. 2A). AAV-F vectors were injected at 10-fold lower dose than AAV9, we have previously demonstrated a greater CNS transduction of AAV-F at a 10-fold lower dose after neonatal ICV delivery(*20*). Furthermore, here in this study, we demonstrated a superior biodistribution in the brain and visceral organs with AAV-F compared to titre matched AAV9 (Fig S4) after neonatal IV delivery. In addition, AAV-F displayed a greater transduction of neurons and astrocytes compared to AAV9 (Fig. S5 & S6).

**Fig. 2.**
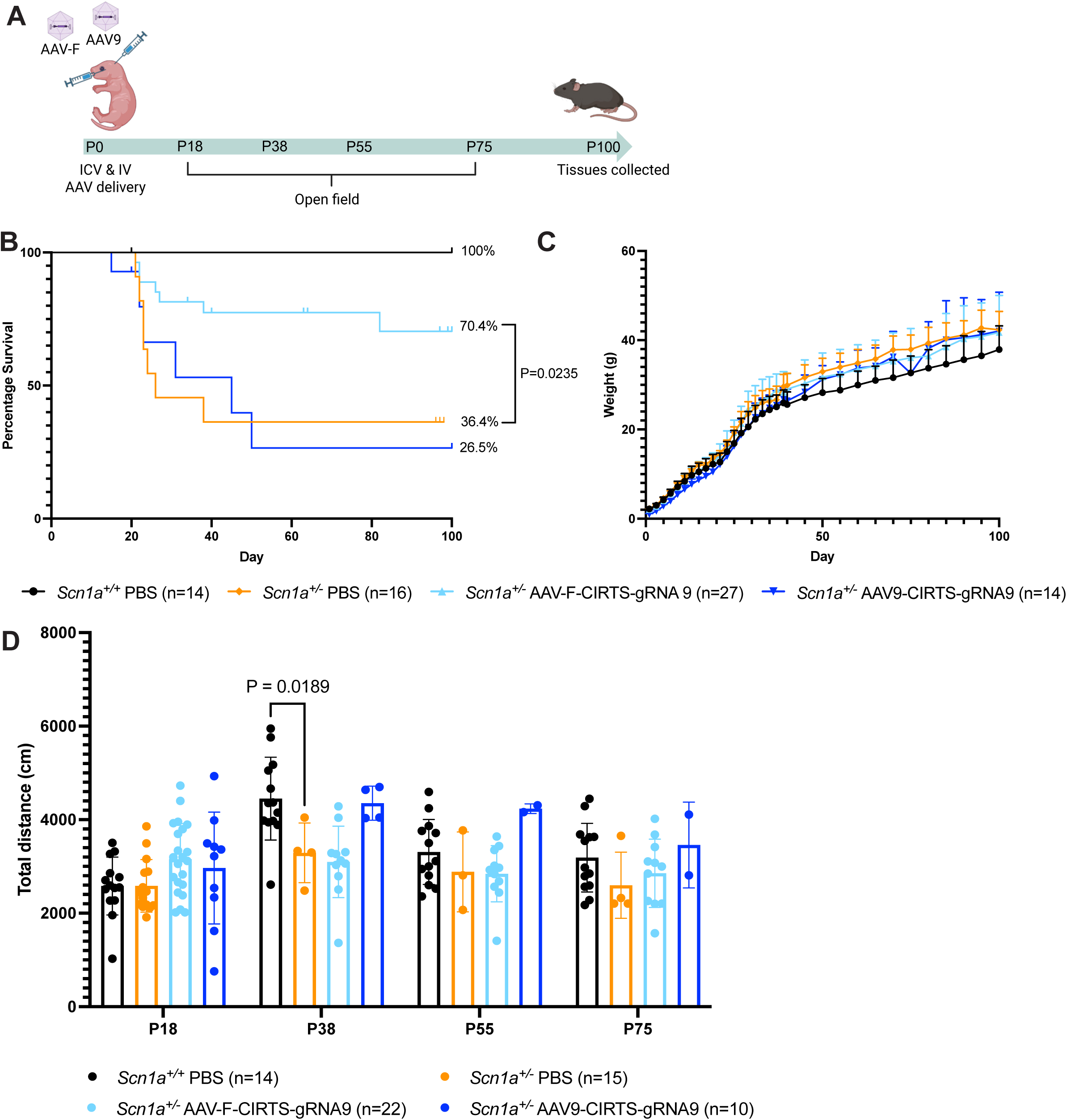
Neonatal AAV-F-CIRTS-gRNA9 RNA editing *in vivo* increases survival of *Scn1a^+/-^* DS mice. *Scn1a^+/-^*mice were administered either AAV-F-CIRTS-gRNA9 (7×10^10^ vg/pup) or AAV9-CIRTS-gRNA9 (7×10^11^ vg/pup) at P0 by intracerebroventricular (ICV) and intravenous (IV) injection, with PBS-treated *Scn1a^+/-^* and *Scn1a^+/+^* littermate controls. (A) Experimental timeline. (B) Kaplan-Meier survival curves of treated mice with percentage survival shown. Log-rank (Mantel-Cox) test. (C) Weights of treated mice. One-way ANOVA with Dunnett’s multiple comparison. (D) Total distance read-outs from open field analyses at 4 different timepoints. Two-way ANOVA with Dunnett’s multiple comparisons.

PBS-injected *Scn1a*^+/+^ and *Scn1a*^+/-^ mice served as controls. *Scn1a*^+/-^ mice treated with AAV-F-CIRTS-gRNA9 had a significant (P=0.0235) increase in overall survival to 70.4% at P100 compared to 36.4% survival at P100 in PBS-treated *Scn1a*^+/-^ controls (Fig. 2A). In contrast, AAV9-CIRTS-gRNA9 only showed an overall survival of 26.5% (P=0.8439, Fig. 2A). The AAV-F-CIRTS-gRNA21 treatment group also appeared less efficient, showing only a 12.5% survival (P=0.0509, Fig. S8A), while AAV9-CIRTS-gRNA21 had a 50% survival at P100 (P=0.3715, Fig. S8A). Weights and open field analysis of neither AAV-F-or AAV9-CIRTS-gRNA9/21 treated mice differed from PBS *Scn1a*^+/+^ and *Scn1a*^+/-^ control group (Fig. 2D, Fig. S7A & Fig. S8C and D). From these data, AAV9 RNA editing at a 10-fold higher titre than AAV-F failed to improve survival we therefore, proceeded with AAV-F-CIRTS-gRNA9 for the remaining experiment.

We interrogated dose-range efficacy of AAV-F-CIRTS-gRNA9 with one higher and one lower dose. High dose (1.75×10^11^ vg/pup) resulted in 65.5% survival (P=0.0732, Fig. 3A), whilst low dose (1.75×10^10^ vg/pup) resulted in 44.4% survival, compared to PBS-treated *Scn1a*^+/-^ mice over 100 days (P=0.5815 and P=0.8556, respectively, Fig. 3A). Neither of the additional AAV-F-CIRTS-gRNA9 doses were superior to our originally tested dose of 7×10^10^ vg/pup. Furthermore, no doses showed difference in weight or open field assessment compared to PBS *Scn1a*^+/+^ control group (Fig. 3B, C and Fig. S9A).

**Fig. 3.**
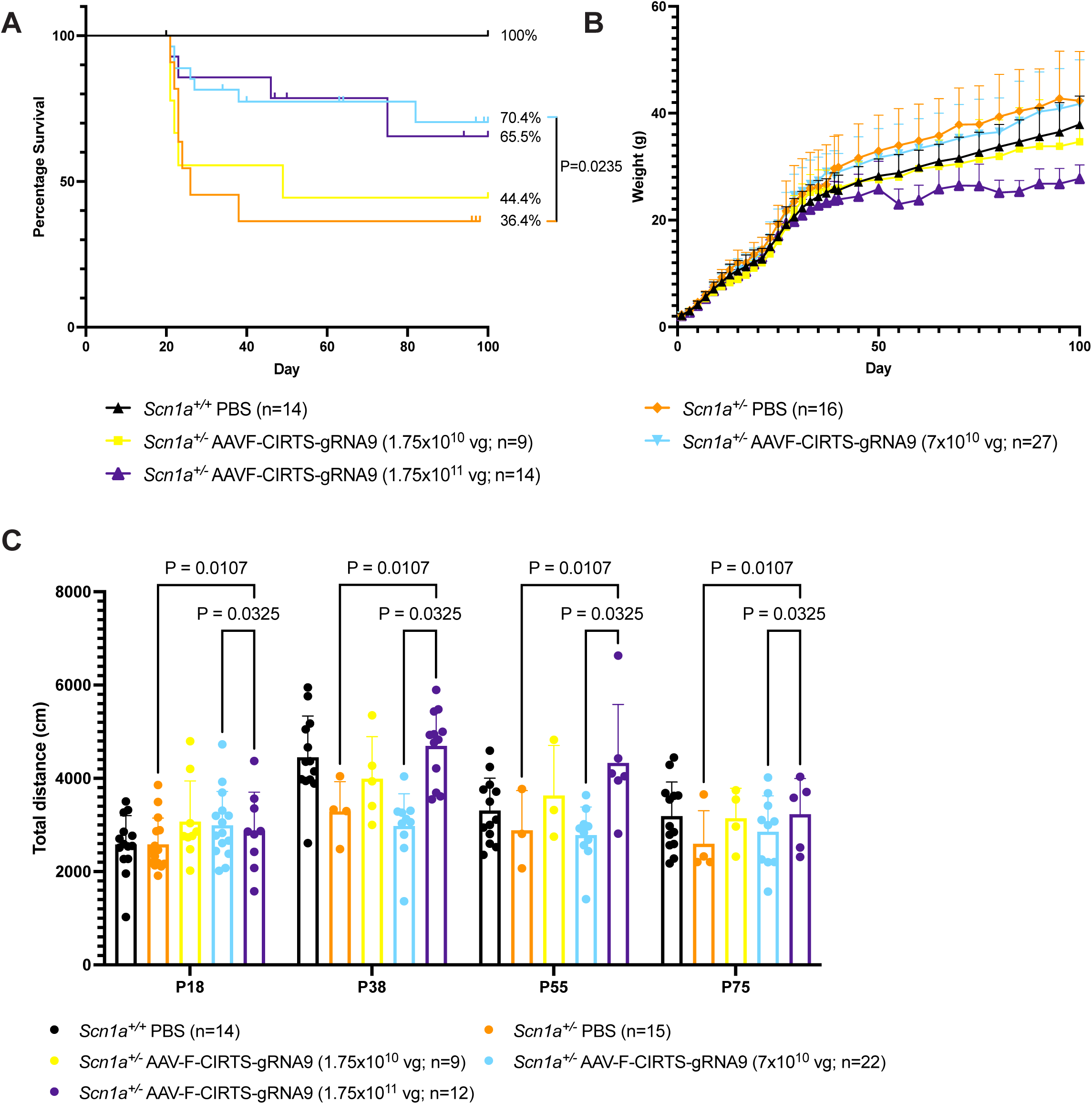
Dosage study of AAV-F-CIRTS-gRNA9 in *Scn1a^+/-^* mice. AAV-F-CIRTS-gRNA9 was administered at 2 additional doses (1.75×10^10^ and 1.75×10^11^ vg/pup) and was compared to the original 7×10^10^ vg/pup cohort presented in Fig. 2. (A) Kaplan-Meier survival curves of treated mice with percentage survival shown. Log-rank (Mantel-Cox) test. (B) Weights of treated mice. AAV-F-CIRTS-gRNA9 1.75×10^11^ vg/pup mice were significantly different from *Scn1a^+/-^* PBS mice at P70 & P80-95 (P=0.0214-0.0451). Mixed-effects analysis with Geisser-Greenhouse correction. (C) Total distance read-outs from open field analyses at 4 different timepoints. Two-way ANOVA with Dunnett’s multiple comparisons.

### AAV-F-CIRTS-gRNA9 reduces seizure burden in DS mice

To assess the effects of this optimal dose of AAV-F-CIRTS-gRNA9 therapy on febrile and spontaneous seizures in *Scn1a*^+/-^ mice, we administered a separate cohort of *Scn1a*^+/-^ mice with neonatal ICV and IV delivery (7×10^10^ vg/pup). We also administered control groups AAV-F-CIRTS-gRNAluc (7×10^10^ vg/pup) and PBS to neonatal *Scn1a*^+/-^ mice via ICV and IV delivery (Fig. 4A). Febrile seizure assessment revealed a significant improvement in seizure severity determined by the Racine scale(*31*) (Fig. 4B), in AAV-F-CIRTS-gRNA9 treated *Scn1a*^+/-^ mice compared to PBS *Scn1a*^+/-^ control group (P<0.0001; Fig. 4B), and AAV-F-CIRTS-gRNAluc *Scn1a*^+/-^ control group (P=0.0111; Fig. 4B).

**Fig. 4.**
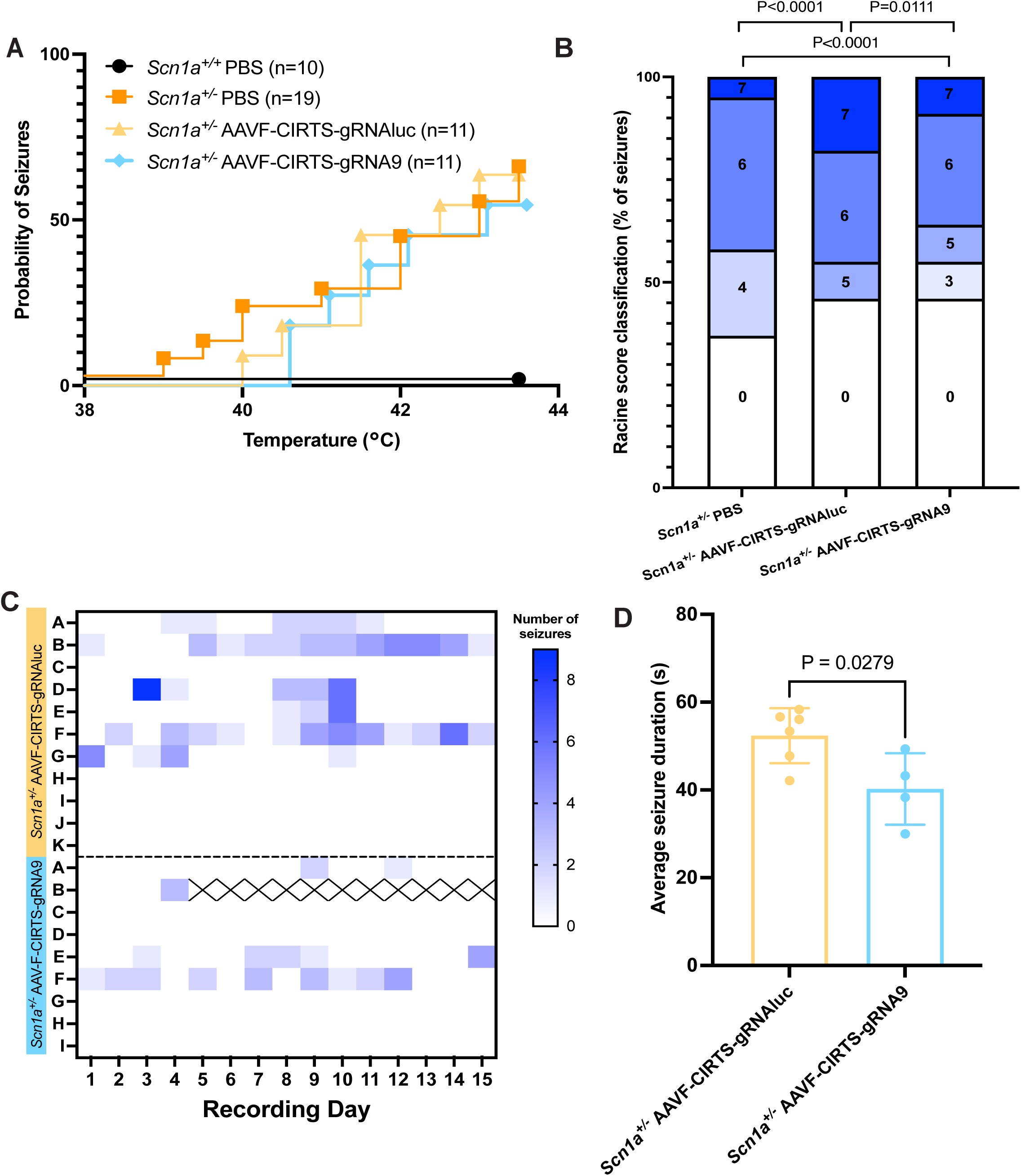
Reduction of seizure burden following AAV-F-CIRTS-gRNA9 treatment. (A) Febrile seizure probability curves. (B) Severity of observed seizures ranked by Racine scale. Fisher’s exact test. (C) Heatmap with coloured squares indicating seizure activity, monitored by EEG. Crosses indicate the death of an animal during EEG recording. (D) Average seizure duration in mice which presented with seizures, during EEG. Unpaired, two-tailed t-test.

Electroencephalogram (EEG) recordings of spontaneous seizures were recorded for 15 days between P30-45 and the results revealed a trend to reduction of seizure burden in AAV-F-CIRTS-gRNA9 mice, compared to a control AAV-F-CIRTS-gRNAluc. Total numbers of seizures (9.7±13 vs 4.1±7.3; P=0.4121; Fig. S10A) and average numbers of seizures per day (0.78±0.93 vs 0.33±0.48; P=0.43; Fig. S10B) were reduced. In the mice which displayed seizures, the average seizure duration was significantly reduced in the treated AAV-F-CIRTS-gRNA9 *Scn1a*^+/-^ group compared to control AAV-F-CIRTS-gRNAluc *Scn1a*^+/-^ group (P=0.0279; Fig. 4D), as well as a trend for reduction in cumulative seizure time (P=0.0640; Fig. S10C).

### AAV-F-CIRTS-gRNA9 normalises molecular signature of DS

To determine the effects of AAV-F-CIRTS-gRNA9 on endogenous *Scn1a* mRNA, we assessed gene expression in the cortex and the heart of treated *Scn1a*^+/-^ mice. Raw wild-type *Scn1a* gene expression levels were restored in both the cortex and heart of AAV-F-CIRTS-gRNA9 mice (P=0.0072 and P=0.0002 vs PBS *Scn1a*^+/-^; Fig. 5A and B).

**Fig. 5.**
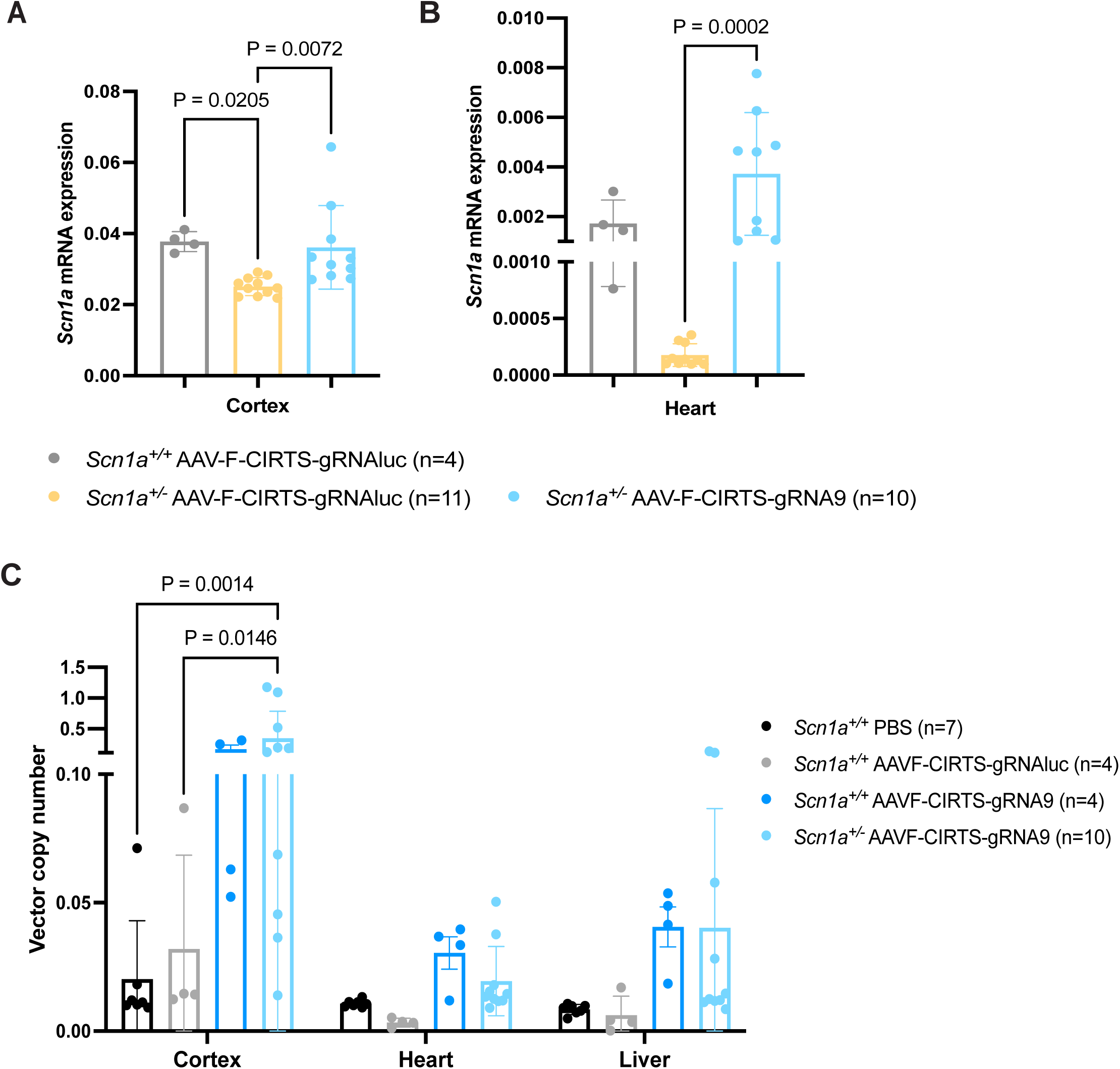
Restoration of *Scn1a* gene expression in the cortex and heart of AAV-F-CIRTS-gRNA9 treated *Scn1a^+/-^* mice. *Scn1a* gene expression in the (A) cortex and (B) heart between *Scn1a^+/+^* and *Scn1a^+/-^* mice (P20-100 tissues) treated with non-targeting AAV-F-CIRTS-gRNAluc vector compared to AAV-F-CIRTS-gRNA9 *Scn1a^+/-^* mice. One-way ANOVA with Dunnett’s multiple comparisons. (C) Vector copy number analysis in P100 cortex, heart and liver. Two-way ANOVA with Tukey’s multiple comparisons.

Vector copy number (VCN) assessing the presence of the effector protein *YTHDF2* genomic sequence, showed an increase in both *Scn1a*^+/-^ and *Scn1a*^+/+^ mice treated with AAV-F-CIRTS-gRNA9 treated mice in the cortex at P100 compared to both PBS (P=0.0014) and AAV-F-CIRTS-gRNAluc (P=0.0146) groups *Scn1a*^+/+^ (VCN 0.35±0.44; Fig. 5C). No difference was observed in the heart and the liver (VCN 0.19±0.14 and 0.04±0.05, respectively; Fig. 5C).

*YTHDF2* mRNA expression mimicked the pattern of vector copy number with highest expression in the cortex in AAV-F-CIRTS-gRNA9 *Scn1a*^+/-^ mice. No significance was observed in the heart and the liver (Fig. S11A).

### *In vivo* safety of AAV-F-CIRTS-gRNA9

Both AAV-F-CIRTS-gRNA9 and AAV-F-CIRTS-gRNAluc were injected to *Scn1a*^+/+^ mice at the two highest dosages 1.75×10^11^ vg/pup and 7×10^10^ vg/pup via neonatal ICV and IV to determine any toxicological findings. No adverse effects on survival or weight were seen (Fig. S12A and B). Open field analyses revealed hyperactivity in the total distance of *Scn1a*^+/+^ AAV-F-CIRTS-gRNAluc mice treated with dose 1.75×10^11^ vg/pup at P18 and P55 (P=0.0003, P=0.0217; Fig. S12C), also with an increase in maximum speed at P55 (P=0.0099; Fig. S12D) compared to PBS *Scn1a*^+/+^ control group.

Immunohistochemical analysis to detect astrocytes (GFAP) and macrophages, including kupffer cells, in the brain and liver respectively was performed post-AAV-F-CIRTS-gRNA9 treatment. No increases in reactive astrocytes were seen in the brains of treated *Scn1a*^+/+^ or *Scn1a*^+/-^ mice (Fig. 6A and B). An increase in the presence of macrophages/kupffer cells in the liver was seen in only the *Scn1a*^+/+^ mice treated with AAV-F-CIRTS-gRNAluc (P=0.0025), this was not observed in mice of either genotype treated with the same dose of AAV-F-CIRTS-gRNA9 (Fig. 6A and C).

**Fig. 6.**
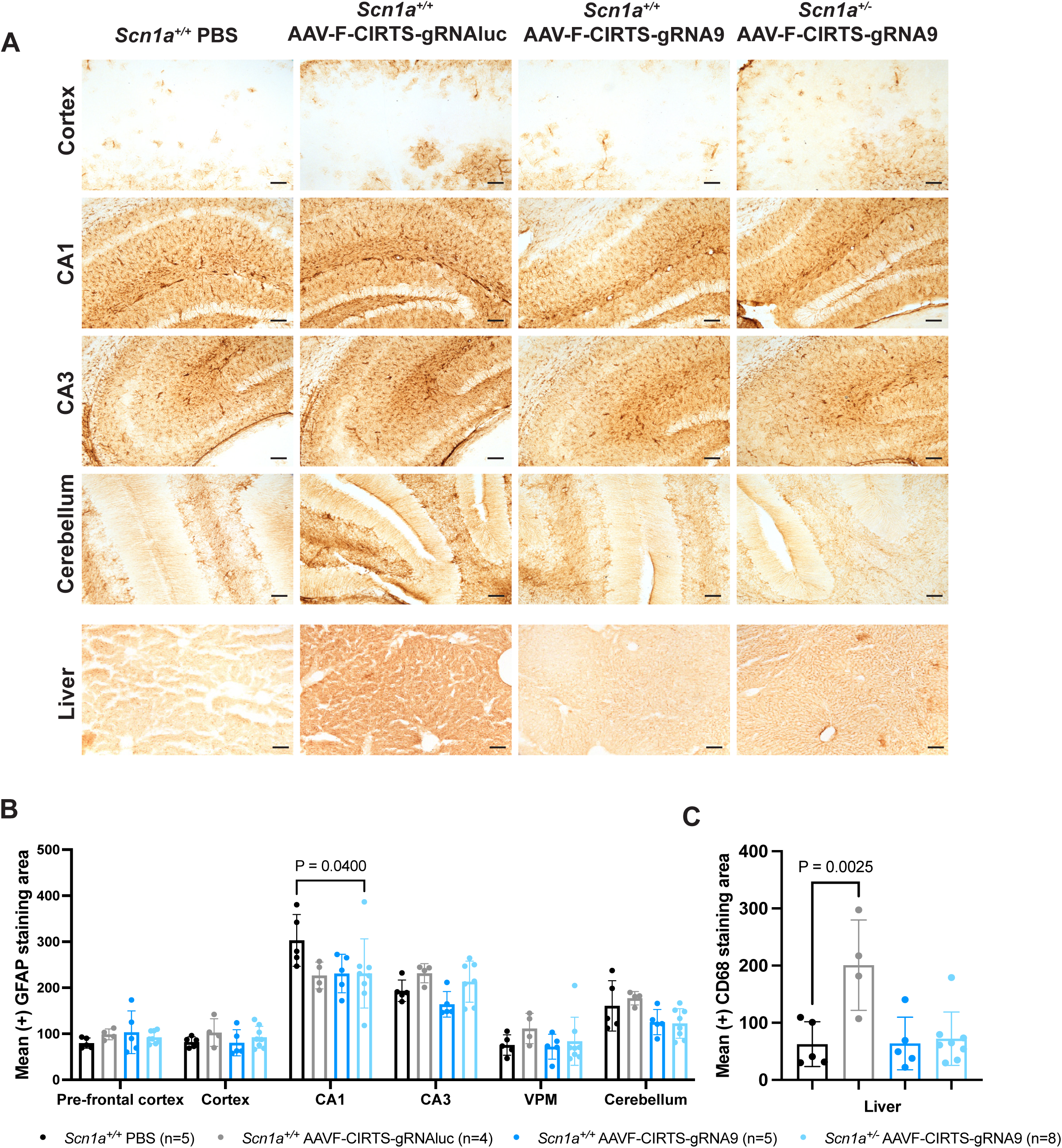
AAV-F-CIRTS-gRNA9 does not lead to toxicity *in vivo*. (A) Representative brain and liver immunohistochemistry images. Scale bar = 100 μm. (B) Quantification of reactive astrocytes (GFAP immunohistochemistry) in the brain of treated mice; CA1 = CA1 region of the hippocampus, CA3 = CA3 region of the hippocampus, VPM = Ventral posteromedial nucleus of the thalamus. Two-way ANOVA with Šídák’s multiple comparisons. (C) Quantification of macrophages (CD68 immunohistochemistry) in the liver of treated mice. One-way ANOVA with Dunnett’s multiple comparisons.

### *In vivo* editing specificity of AAV-F-CIRTS-gRNA9

To identify gRNA-specific changes in gene expression, we conducted RNA-seq on cortical tissue 50 days post-injection and compared AAV-F-CIRTS-gRNAluc and AAV-F-CIRTS-gRNA9 *Scn1a*^+/-^ mice. Only 9 differentially expressed genes were found; 3 up-regulated and 6 down-regulated (Fig. 7A and B). Of note, inflammation-associated genes were down-regulated including *Igkc* (−7.27Log2FC, P=0.0417), *Cxcl13* (−7.35Log2FC, P=0.0417) and *Adgre* (−1.86Log2FC, P=0.0019) and the only driver GO term changed across the comparison, was the down-regulation of CXCR5 chemokine receptor binding (Fig. 7C; P=0.0499). None of the *Scna* genes, which encode other voltage-gated sodium ion channels, were differentially expressed in the gRNA9 group compared to gRNAluc.

**Fig. 7.**
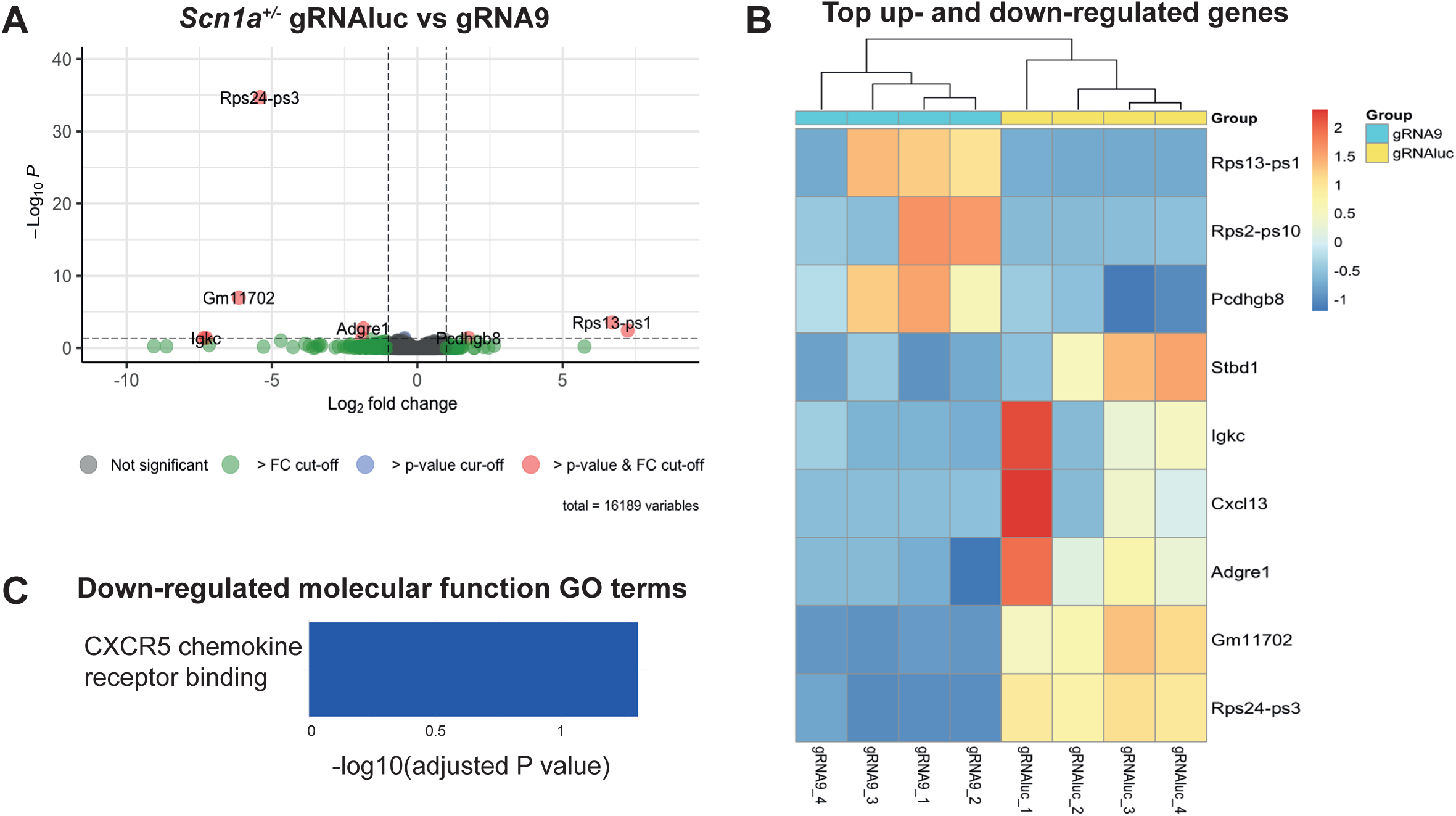
RNA-seq depicting differential gene expression following AAV-F-CIRTS-gRNA9 treatment. *Scn1a^+/-^* cortical tissue analysed at P50. Comparison of AAV-F-CIRTS-gRNAluc and AAV-F-CIRTS-gRNA9 mice. (A) Volcano plot showing the main differentially expressed genes (DEGs). (B) Heatmap showing top up-and down-regulated genes per comparison, presented as z-score, ranked by P value. Driver gene ontology (GO) term classification of the molecular function of (C) down-regulated DEGs. No molecular function GO terms for up-regulated genes were identified.

## DISCUSSION

Here, we provide preclinical proof-of-concept data demonstrating that CIRTS-mediated RNA editing can mitigate the phenotype of a severe neurological disorder, Dravet syndrome. A single administration of AAV-F-CIRTS-gRNA9 by combined ICV and IV delivery to *Scn1a^+/-^* DS mice induces destruction of *Scn1anat* and completely rescues endogenous *Scn1a* mRNA expression in both the cortex and the heart. This was shown to improve disease manifestation by reducing febrile seizure severity and significantly reducing SUDEP, with minimal off-target effects; whilst being compromised entirely of human components.

We designed the gRNAs used in this study with the aim of restoring wild-type *Scn1a* expression levels, without leading to supraphysiological expression. Notably, modest 2.6-and 1.9-fold increase of *Scn1a* expression *in vitro* were achieved with gRNA21 and gRNA9 respectively. We sought to combine these with AAV-F, an engineered capsid, shown to have a more efficient CNS transduction profile than AAV9 in rodents(*32*), human neurons(*32*) and NHPs(*19*). Furthermore, we have validated this with neonatal delivery both by ICV(*20*) and IV delivery. By packaging AAV-CIRTS-gRNA9 into AAV-F and AAV9 we demonstrated vastly different therapeutic effects with 70.4% and 26.5% survival, respectively, strikingly this was achieved with AAV-F being administered at a 10-fold lower dose. However, gRNA21 delivered by either AAV-F or AAV9 did not reveal a significant improvement in survival. We tested a varying dose of AAV-F-CIRTS-gRNA9 and determined that 7×10^10^ vg/pup achieved a significant survival of 70.4% at P100, interestingly increasing the dose further to 1.75×10^11^ vg/pups showed a reduced survival of 65.5% over 100 days. These findings suggest a narrow therapeutic dose ranging window for a successful treatment for DS. This is particularly important in the context of *SCN1A*, as gain-of-function *SCN1A* variants, which are known to cause familial hemiplegic migraine type 3 and a severe infantile developmental and encephalopathy (distinct from DS) with associated movement disorder(*33*). Therefore, future studies require a more refined dose ranging evaluation of AAV-F with gRNA9 or gRNA21 to determine the optimal therapeutic dose, with appropriate levels of physiological *Scn1a* and no behavioural side effects.

As a main symptom of DS is seizures, we sought to assess the seizure activity in our AAV-F-CIRTS-gRNA9 treated *Scn1a^+/-^* DS mice via febrile seizure assessment and EEG monitoring for 15 days (P30-45). We delivered a vehicle and control vector AAV-F-CIRTS-gRNAluc to *Scn1a^+/-^*DS mice as a comparators. Febrile seizure assessment revealed a significant reduction in seizure severity determined by the Racine scale in AAV-F-CIRTS-gRNA9 treated *Scn1a^+/-^*mice compared to PBS and AAV-F-CIRTS-gRNAluc *Scn1a^+/-^* control group. EEG seizure monitoring was performed for AAV-F-CIRTS-gRNA9 and AAV-F-CIRTS-gRNAluc treated *Scn1a^+/-^*DS mice. Our results showed that we could significantly reduce the seizure duration in our treated *Scn1a^+/-^* DS mice. From these data, we observed a variability in the AAV-F-CIRTS-gRNAluc *Scn1a^+/-^* DS mice, as some of these mice did not display seizures during the EEG recording. Which may partly be due to the variable penetrance of spontaneous seizures in this model(*34*). This was a limitation in our study and future studies, will require further power calculations to establish the appropriate number of *Scn1a^+/-^* mice required for EEG seizure monitoring.

Molecular assessment revealed a significant increase in endogenous *Scn1a* mRNA in the cortex and the heart of treated AAV-F-CIRTS-gRNA9 *Scn1a^+/-^* mice. Immunohistochemical analysis in the brain of AAV-F-CIRTS-gRNA9 *Scn1a^+/-^* and *Scn1a^+/+^* mice showed no signs of toxicity. Liver toxicity was only observed in AAV-F-CIRTS-gRNAluc *Scn1a^+/+^* treated group, suggesting potential guide-specific side-effects of AAV-F-CIRTS-gRNAluc vector.

To determine the off-target effects of our gRNA9, we conducted RNA-seq on cortical brain tissues for AAV-F-CIRTS-gRNAluc and AAV-F-CIRTS-gRNA9 *Scn1a*^+/-^ treated mice. The data revealed inflammation-associated genes including *Igkc*(*35*), *Cxcl13*(*36*) and *Adgre*(*37*) were down-regulated in the AAV-F-CIRTS-gRNA9 treated group. Previously, up-regulation of the CXCL13-CXCR5 signalling pathway has been associated with intractable epilepsy(*36*), suggesting that our therapy may be able to reduce some molecular signatures of epilepsy pathogenesis. Furthermore, our data suggest no off-target effects of the gRNA9 coupled with the CIRTS machinery.

Several approaches in preclinical development have used genetic therapies specifically in inhibitory interneurons using *SCN1A* gene replacement with split-intein dual AAVs(*38*) and Adenoviral vectors(*39*), CRISPR/dCas9 activation of *SCN1A*(*40, 41*) and transcription factor-mediated *SCN1A* up-regulation(*30*). While these approaches target the primary pathology of Dravet syndrome (DS), our strategy complements these alternative therapeutic interventions by also considering effects in less well-studied cell populations, such as cardiomyocytes(*10*). Here, we have shown that our AAV-F-CIRTS-gRNA9 can increase *Scn1a* mRNA in the cortex and the heart.

*SCN1ANAT* has been detected in surgical brain tissue specimens from patients (1-19 years of age) undergoing resection to treat refractory, non-DS epilepsy(*42*). Previous evidence has shown that targeting *Scn1anat* using small oligonucleotides termed AntagoNATs were able to reduce seizures and normalise neuronal activity in DS knock-in mice(*27*). Furthermore, we have also shown that these AntagoNATs can be vectorised with AAV9, which also demonstrates a significant improvement in DS phenotype(*43*). Here, we also are targeting *Scn1anat*, but instead of blocking the interaction between this and the *Scn1a* mRNA, we are inducing the degradation of *Scn1anat*, with similar phenotypic improvement. These findings demonstrate the feasibility of using AAV mediated RNA editing to modulate *Scn1a* expression, highlighting its potential as a therapeutic for DS.

Genetic therapies for DS have progressed to clinical evaluation. These include the antisense oligonucleotide (ASO) zorevunersen, currently under investigation in Phase 1/2a MONARCH and ADMIRAL trials, which involve repeated intrathecal administration(*44*). Encoded Therapeutics has developed an AAV9-based transcription factor(*30*) approach specifically targeting GABAergic interneurons, now being assessed in a Phase 1/2 clinical trial utilizing ICV delivery alone(*45*). Similar to Encoded Therapeutics’ strategy, our AAV-F-CIRTS-gRNA9 platform offers a one-time treatment; however, it confers the additional advantage of enabling broader targeting of CNS cell populations expressing Nav1.1. Furthermore, the inclusion of IV administration is intended to achieve efficient cardiac transduction, addressing evidence implicating altered cardiac electrophysiology(*8, 11*) in contributing to SUDEP.

Our study presented limitations. Some of the *Scn1a^+/-^* DS mice did not present seizures during EEG recording and therefore, for future studies we will require further power calculations. The future studies required have been detailed in this discussion.

In conclusion, we have developed and characterized an AAV-F-CIRTS RNA editing therapeutic strategy for the treatment of DS. Incorporation of gRNA targeting *Scn1anat* successfully modulated *Scn1a* expression and significantly improved survival, seizure severity and duration in *Scn1a*^+/-^ mice. These findings provide compelling proof-of-concept evidence supporting the potential of this approach as a therapeutic intervention for DS.

## MATERIALS AND METHODS

### Study design

This study was designed to investigate if AAV-CIRTS-mediated RNA editing to induce the destruction of *Scn1anat* could lead to therapeutic benefit in a clinically relevant DS mouse model. The following design elements were included in this study. N=3 independent experiments were conducted for *in vitro* gRNA screening experiments with at least n=3 technical replicates per condition. Mice were assigned randomly to study groups, with at least one control (either PBS or AAV-F-CIRTS-gRNAluc-treated) *Scn1a*^+/-^ mouse per litter and remaining *Scn1a*^+/+^ and *Scn1a*^+/-^ littermates mice treated with AAV-F-CIRTS-gRNA9/21 or AAV9-CIRTS-gRNA9/21 treated mouse. Prospectively selected endpoints of these studies were survival, weight, open field, longitudinal gene expression and neuropathological analyses. Sample sizes for *in vivo* experiments were not pre-defined before the outset of data collection as no preliminary data were available to estimate effect sizes and thus be able to complete power analyses. *In vivo* studies analysing seizures used *Scn1a*^+/-^ littermates treated with AAV-F-CIRTS-gRNAluc as a non-targeting guide as a control for AAV-F-CIRTS-gRNA9. At termination of the in-life period for all *in vivo* experiments, mice were assigned a numeric code and investigators were blinded to treatment and genotype whilst analysing tissue.

### gRNA design and cloning

The *Scn1anat* target sequence was inputted into sOligo software (https://sfold.wadsworth.org/cgi-bin/soligo.pl), varying the parameter for gRNA length. Folding temperature was set to 37°C and ionic conditions set to 1M NaCl with no divalent ions. Resulting outputs were screened for those with less than -8.0 Gibbs free energy with positional clusters identified. Potential gRNA sequences were manually mapped to RNA secondary structures produced using mFold software (http://www.unafold.org/mfold/applications/rna-folding-form.php) to determine qualities of each gRNA such as the number of termini and percentage of gRNA sequence in open conformation. gRNAs were cloned into the AAV-CIRTS plasmid (Addgene plasmid #132544), using gBlocks from Integrated DNA Technologies and InFusion cloning (Takara Bio). The AAV-CIRTS plasmid contains a cytomegalovirus promoter (CMV), driving ssRNA binding protein β-defensin 3, RNA hairpin-binding protein (TBP6.7), effector protein (YTHDF2) and U6 promoter driving gRNA (referred to as AAV-CIRTS in the main text, Fig. 1B).

### In vitro analyses

For luciferase assays; HEK293T cells (5×10^4^ per well of 24-well plate) were transfected with 0.8μg of an AAV plasmid containing an Spleen focus forming virus ubiquitous promoter (SFFV) driving luciferase linked to green fluorescent reporter gene with a T2A linker peptide(*46*) (referred to as AAV-SFFV-*luciferase*-T2A-*eGFP* in the main text) *±* 0.8μg AAV-CIRTS-gRNAluc using Lipofectamine 2000 (Thermo Fisher Scientific). Cells were lysed in Luciferase Cell Culture Lysis Reagent (Promega) at 72h post-transfection, with half the lysate used to determine protein concentration using the DC protein assay (BioRad). The remaining half of the lysate was used for luminometry measured in a FLUOstar® Omega microplate reader (BMG Labtech). Luciferase expression was normalised to protein concentration within each sample.

For *Scn1anat* gRNA screening; Neuro2a cells (3×10^4^ per well of 24-well plate) were differentiated in DMEM (Thermo Fisher Scientific), 2% FCS (Sigma-Aldrich), 0.5mM cAMP (Merck) and 20μM Retinoic Acid (Merck). At 6DIV, cells were transfected with 0.8μg AAV-SFFV-*luciferase*-T2A-*eGFP* or AAV-CIRTS-gRNA1-21 using Lipofectamine 2000 (Thermo Fisher Scientific). Cells were lysed in TRIzol reagent (Thermo Fisher Scientific) at 5d post-transfection. RNA was extracted and qPCR used to determine changes in *Scn1a* gene expression (protocols described below).

### AAV vector production

Recombinant AAV vectors were produced by triple-transfection(*47*) of AAVpro 293T cells (Takara Bio) with AAV-CIRTS, adenovirus helper plasmid (Harvard University) and either AAV9 (University of Pennsylvania; Addgene #112865) or AAV-F (Harvard University, Addgene #166921) capsid plasmids using PEImax (Polysciences Inc.). HPLC using POROS CaptureSelect AAVX Resin (Thermo Fisher Scientific) was used to purify resulting vector before being titrated by qPCR.

### Animals

All procedures were performed in accordance with UK Home Office Animals (Scientific Procedures) Act 1986 under PPLs 70/14300, PP8023218 & PP9223137. All mice were housed in IVCs with enrichment, under a 12 hour:12 hour light–dark cycle and had access to food and water *ad libitum*. The mouse model of Dravet syndrome used in this project is derived from the 129Sv-*Scn1a^tm1Kea^*/Mmjax model(*28, 29*) (Jackson Laboratory) that has been extensively used in DS pre-clinical research. Males from this strain were bred with wild-type CD1 females to obtain a F1 129Sv-*Scn1a^tm1Kea^*/Mmjax x CD1 generation. F1xF1 crosses were bred to obtain F2 129Sv-*Scn1a^tm1Kea^*/Mmjax x CD1 heterozygote females which were bred to wild-type C57BL/6J males to obtain experimental litters (here referred to as *Scn1a*^+/-^ in the text). Humane endpoints of >15% weight loss, observation of two seizures and/or clear signs of illness (piloerection, hunched posture and laboured breathing).

At post-natal day 0 (P0), *Scn1a*^+/-^ pups were injected by bilateral intracerebroventricular (ICV; 5 μL per hemisphere, using previously published coordinates(*48*)) and intravenous (IV; 25 μL via superficial temporal vein(*49*)) routes of administration with either PBS, AAV9 (total dose 7×10^11^ vg/pup (2×10^11^ vg ICV + 5×10^11^ vg IV)) or AAV-F (total dose 1.75×10^10^ vg/pup (5×10^9^ vg ICV + 1.25×10^10^ vg IV), 7×10^10^ vg/pup (2×10^10^ vg ICV + 5×10^10^ vg IV) and 1.75×10^11^ vg/pup (5×10^10^ vg ICV + 1.25×10^11^ vg IV)) using a 33-gauge Hamilton needle (VWR). For the biodistribution study AAV9 (1.9×10^10^ and 1.9×10^11^ vg/pup) and AAV-F (1.9×10^10^ vg/pup) was delivered intravenously via superficial temporal vein(*49*)). After the procedure, mice were returned to their dams. Weight and survival were monitored until endpoint.

Open field analysis with 15 minutes free locomotion was completed at P18, P38, P55 and P75 using SMART v3 software (Harvard Panlab). At endpoint, tissues were collected by snap-freezing or fixing in 4% PFA.

### EEG recordings

AAV-F-CIRTS-gRNAluc and AAV-F-CIRTS-gRNA9 treated *Scn1a*^+/-^ mice underwent EEG recordings from P30-P45 using wireless electrocorticogram (ECoG) transmitters (Cat. No. A3048P2-AA-C37-D, single-channel transmitter, Open Source Instruments), as previously described(*43*).

### RNA/DNA extraction, gene expression and VCN

RNA extraction was completed using the PureLink kit (Thermo Fisher Scientfic), DNA extraction was completed using DNeasy Blood & Tissue kit (Qiagen), as previously described(*43*). The following primers and probes were used to quantify gene expression, titrate AAV vectors and perform vector copy number analyses. *Scn1a;* forward: TCAGAGGGAAGCACAGTAGAC, reverse: TTCCACGCTGATTTGACAGCA, probe: CCAGAAGAAACCCTTGAGCCCGAA (fluorophore: ABY, quencher: QSY, Thermo Fisher Scientific). *Gapdh*; forward: ACGGCAAATTCAACGGCAC, reverse: TAGTGGGGTCTCGCTCCTGG, probe: TTGTCATCAACGGGAAGCCCATCA (fluorophore: VIC, quencher: QSY, Thermo Fisher Scientific). *YTHDF2*; forward: CCTCCATTGGCTTCTCCTATTC, reverse: GTTGCTCAGCTGTCCATAAGA, probe: TTAAGTAGGGCATGGCTGTGTCACC (fluorophore: FAM, quencher: ZEN / Iowa Black™ FQ, Integrated DNA Technologies).

The same sequence *Gapdh* primers and probe, and *Scn1a* primers were used for ddPCR analysis, but with the *Scn1a* probe fluorophore was adapted from ABY to FAM. 5µL of each cDNA (equivalent to 85 pg) was loaded per reaction with 1X ddPCR Multiplex Supermix (Bio-Rad) and 4 mM DTT. ddPCR analysis was performed using the BioRad QXOne in the UCL NeuroGTx Vector Core Facility. The Facility is supported by the NIHR Great Ormond Street Hospital Biomedical Research Centre. The views expressed are those of the author(s) and not necessarily those of the NHS, the NIHR or the Department of Health.

### Immunoperoxide staining

40 μm coronal sections cut using a Micron HM 430 freezing microtome (Thermo Fisher Scientific) were analysed using previously published protocols(*43*). Neuropathological analyses used the following primary antibodies; anti-GFAP (Proteintech 16825-1-AP; 1:2000) and anti-CD68 (BioRad MCA1957; 1:100). Images were acquired using a Leica Microsystems stereoscopic microscope (DM4000), Leica DFC7000T camera and Leica LAX X software (all Leica Microsystems UK). Images were background corrected using FIJI (ImageJ) and thresholding software (Image-Pro Premier v10; Media Cybernetics) was used to quantify expression.

### Immunofluorescence staining

40 μm coronal sections were used(*43*). Immunolabeling was performed on free-floating sections using antibodies against GFP (Abcam; ab290), NeuN (Merek; MAB377) and GFAP (Proteintech; 16825-1-AP)(*20*). Images were assessed using fluorescence microscope (Leica DFC7000 7) using LAS X Microscope Software (Leica Microsystems). The percentage of GFP positive neurons and astrocytes were analysed using the CellProfiler Software (Broad Institute 2021). The images were overlayed using ImageJ.

### RNA-seq

Total RNA was extracted from the cortex of *Scn1a^+/+^* and *Scn1a^+/-^* P50 mice that were controls, or had been treated with, AAV-F-CIRTS-gRNAluc or AAV-F-CIRTS-gRNA9.

Untreated control samples were processed by UCL Genomics using a KAPA mRNA HyperPrep and NextSeq2000 P2 100c configuration with single-end reads at 25M reads/sample. N=8 mice per genotype were submitted, but due to a pooling error, *Scn1a^+/+^*Untreated_5 could not be included in the analysis.

AAV-CIRTS-treated samples (n=4 per group) were processed by Azenta Life Sciences using Illumina NovaSeq with paired-end reads in 2×150 bp configuration. Trimmomatic v.0.36 was used to trim sequence reads to remove adapter sequences before reads were mapped to *Mus musculus* GRCm38 reference genome using the STAR aligner v.2.5.2b. Unique gene counts were extracted using featureCounts (Subread package v.1.5.2).

Both RNA-seq analyses then used DESeq2(*50*) to determine differential gene expression analysis. Untreated samples were subjected to an absolute Log2 fold change > 0.7, whilst AAV-CIRTS comparisons used an absolute Log2 fold change > 1. In both analyses, genes with an adjusted P value < 0.05 were called differentially expressed genes. A gene ontology analysis was performed on the statistically significant set of genes by implementing enrichGO and clusterProfiler R packages(*51*) to cluster the set of genes based on their molecular function and determine their statistical significance.

### Statistical analyses

GraphPad Prism (version 10.3.1, Boston) was used for statistical analyses. All continuous variables are presented as mean±SD in graphs. *In vitro* data were analysed using an unpaired, two-tailed t-test for luciferase assay or one-way ANOVA with Dunnett’s multiple comparisons for gRNA screening. Survival was determined using a Log-rank (Mantel-Cox) test. Weights were initially screened for differences with one-way ANOVA with Dunnett’s multiple comparisons for group-group comparisons and a mixed-effect analysis was completed with analysis at each individual timepoint. Groups were excluded in mixed-effect analyses from the timepoint that only a single mouse remained. Open field was analysed using two-way ANOVA with Dunnett’s multiple comparisons. Racine scaling of febrile seizures were contingency analysed with Fisher’s exact test. A mixed-effects analysis was used to compare overall EEG seizure burden, with either Mann-Whitney or unpaired, two-tailed t-test for individual seizure readouts. One-way and two-way ANOVAs were used to analyse all molecular data.

## Acknowledgments

We would like to thank A. Rahim and J. Baruteau for the use of their Home Office PPL. We would like to thank, R.Towns, M. Bandol, N. Foster, N. Marzouki and the staff from Biological Services for their help with maintenance of the DS colony at University College London. Figures 1A and 2A were created in BioRender. Antinao Diaz, J. (2025) https://BioRender.com/k81r069.

## Author contributions

Conceptualization: R.K and S.N.W. Methodology: R.K, M.M and E.M.C. Investigation: R.K, E.M.C, J.F.A.D, A.K, Z.W, V.F, A.A.B, G.L. Visualization: R.K and E.M.C. Funding acquisition: R.K, S.S, M.M, S.N.W and G.L. Project administration: R.K. Writing – original draft: R.K and E.M.C. Writing – review & editing: all authors.

## Data and materials availability

All data associated with this study are present in the paper in Supplementary Material. Requests for data should be addressed to R.K.

---

**Fig. S1.**
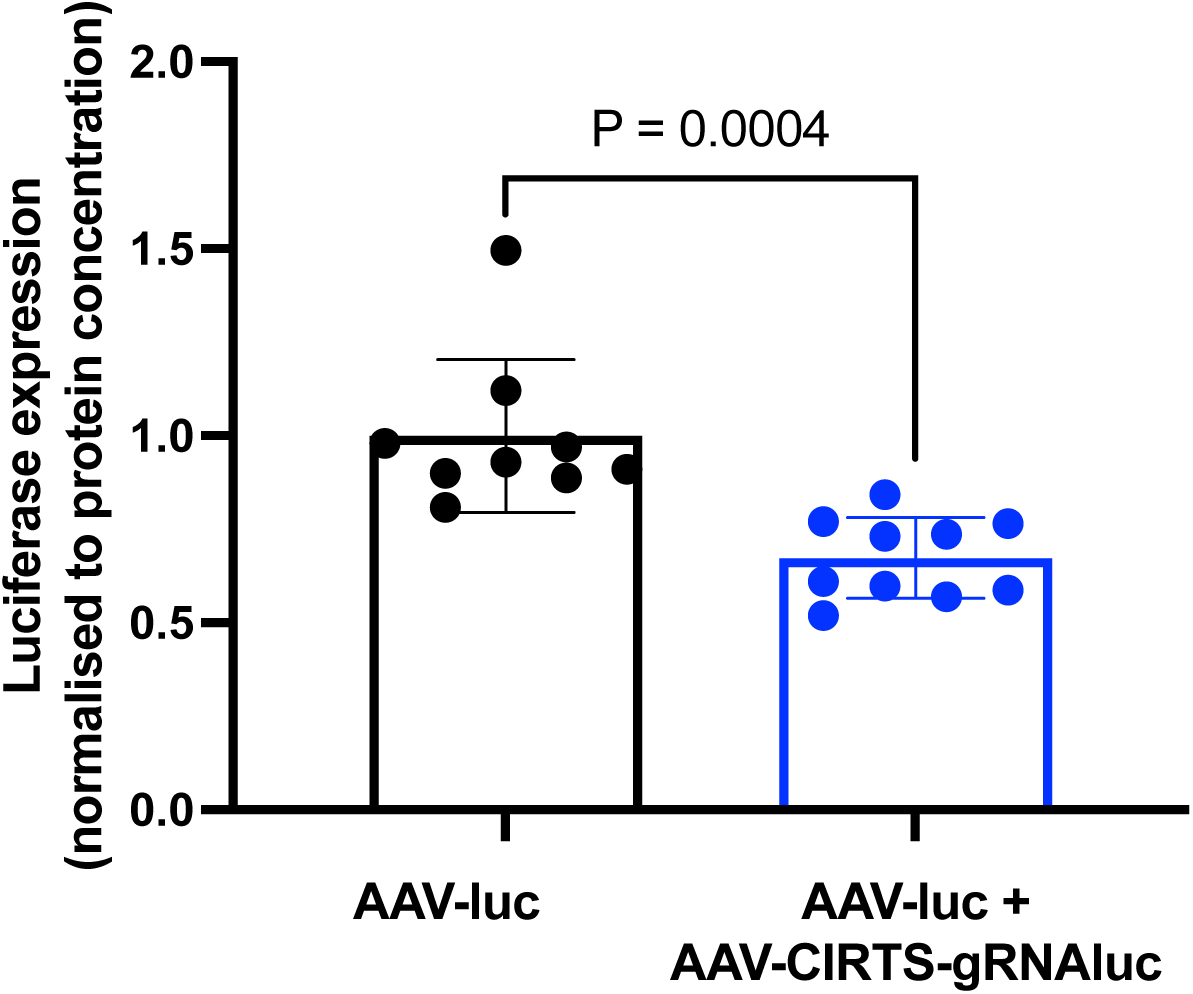
*In vitro* validation of AAV-CIRTS machinery. AAV-CIRTS-gRNAluc significantly down-regulated luciferase expression in HEK293T cells after transient co-transfection with AAV-SFFV-*luciferase*-T2A-*eGFP* plasmid. Luciferase expression normalised to protein concentration within lysate. Unpaired, two-tailed t-test.

**Fig. S2.**
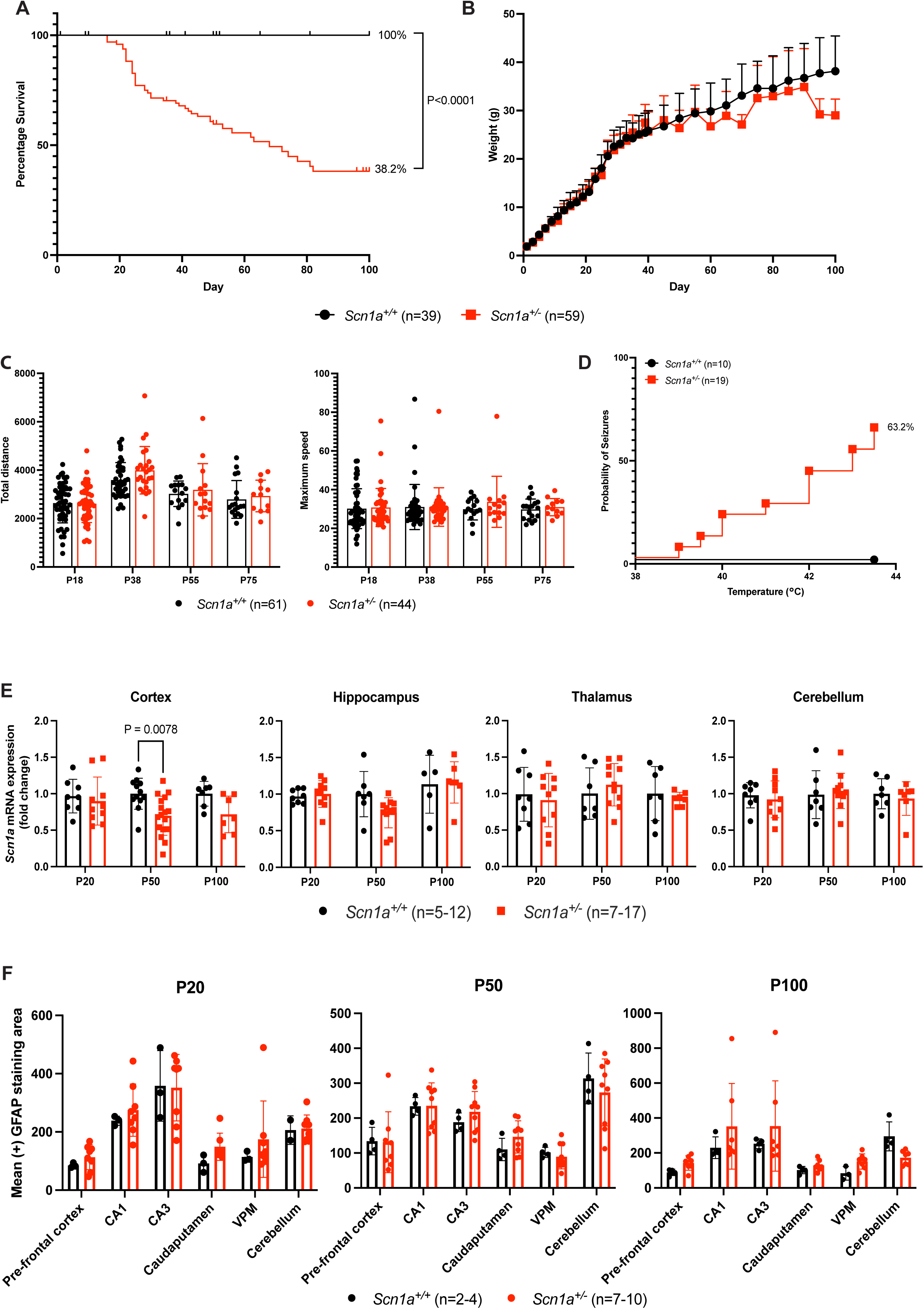
Characterisation of untreated *Scn1a^+/-^* DS mice. (A) Kaplan-Meier survival curves of untreated mice over 100 days of development with percentage survival shown. Log-rank (Mantel-Cox) test. (B) Weights of untreated mice. (C) Total distance and maximum speed read-outs from open field analyses at 4 different timepoints. Two-way ANOVA with Dunnett’s multiple comparisons. (D) Induction of febrile seizures with temperature of seizure onset. (E) *Scn1a* gene expression in 4 brain regions. Two-way ANOVA with Šídák’s multiple comparisons within each brain region. (F) Quantification of reactive astrogliosis (GFAP immunohistochemistry) in the brain of untreated mice; CA1 = CA1 region of the hippocampus, CA3 = CA3 region of the hippocampus, VPM = Ventral posteromedial nucleus of the thalamus.

**Fig. S3.**
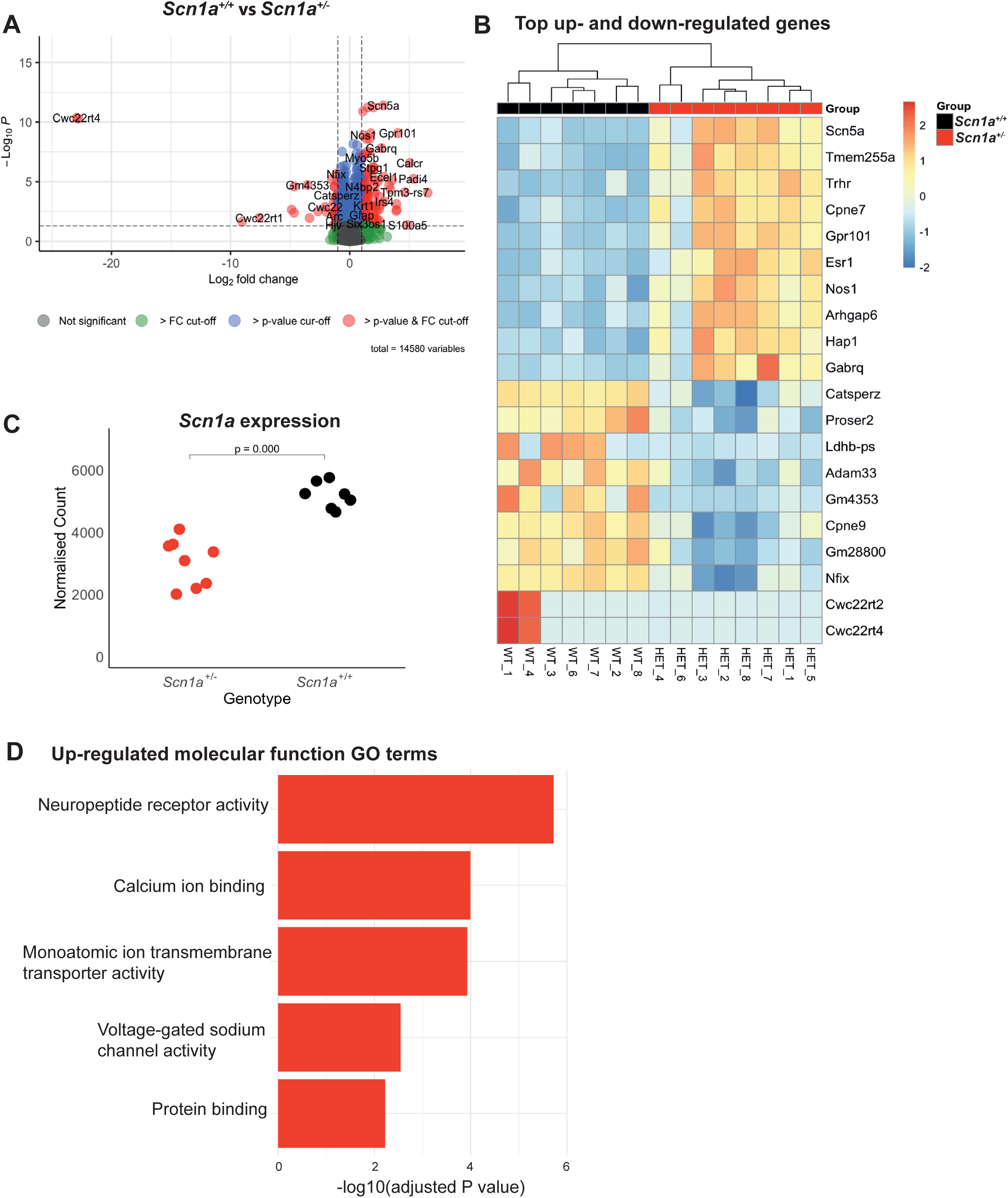
RNA-Seq analysis of untreated *Scn1a^+/-^* DS mice. *Scn1a^+/+^*and *Scn1a^+/-^* cortical tissue analysed at P50. (A) Volcano plot showing the main differentially expressed genes. (B) Heatmap showing top 10 up-and down-regulated genes, presented as z-score, ranked by P value. (C) Confirmation of *Scn1a* down-regulation in *Scn1a^+/-^* mice with normalised read count, (−0.77Log2FC P=5.85×10^-6^). (D) Driver gene ontology (GO) term classification of the molecular function of up-regulated DEGs. No molecular function GO terms for down-regulated genes were identified.

**Fig. S4.**
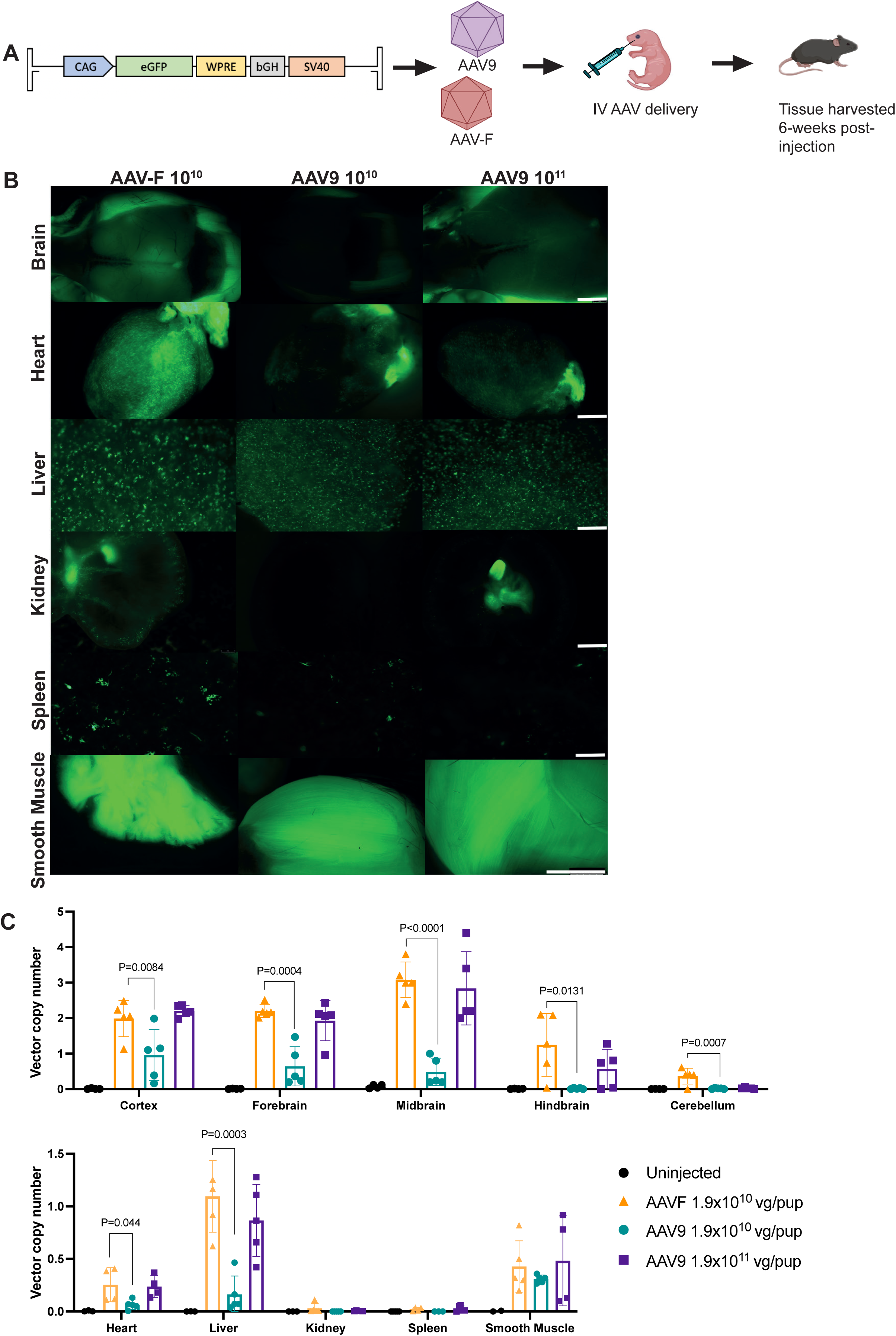
Increased biodistribution and transgene expression of AAV-F-CAG-GFP vector compared to AAV9. Wild-type C57BL/6J mice were administered either AAV-F-CAG-GFP (1.9×10^10^ vg/pup) or AAV9-CAG-GFP (1.9×10^10^ vg/pup or 1.9×10^11^ vg/pup) at P0 by intravenous (IV) injection, with uninjected littermate controls. (A) Experimental timeline. (B) *Ex vivo* images of organs from injected mice. Scale bar = 2mm. (C) Quantification of vector copy number in discreet brain regions and visceral organs. One-way ANOVA with Dunnett’s multiple comparisons.

**Fig. S5.**
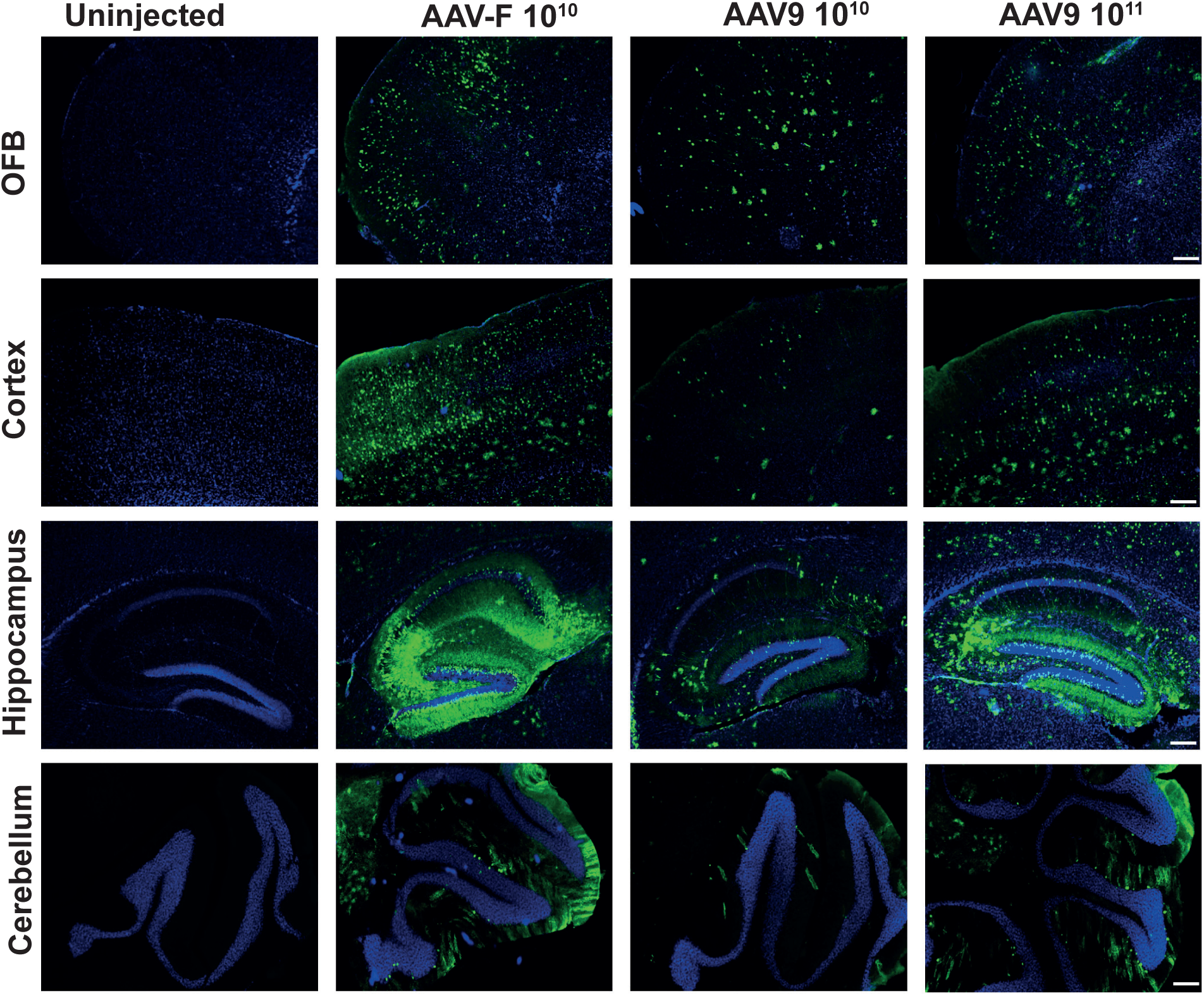
GFP biodistribution in the brain of AAV-F and AAV9-CAF-GFP vectors. Immunofluorescence stain showing GFP positive cells in the olfactory bulb (OFB), cortex, hippocampus and cerebellum. Scale bar shown on the images.

**Fig. S6.**
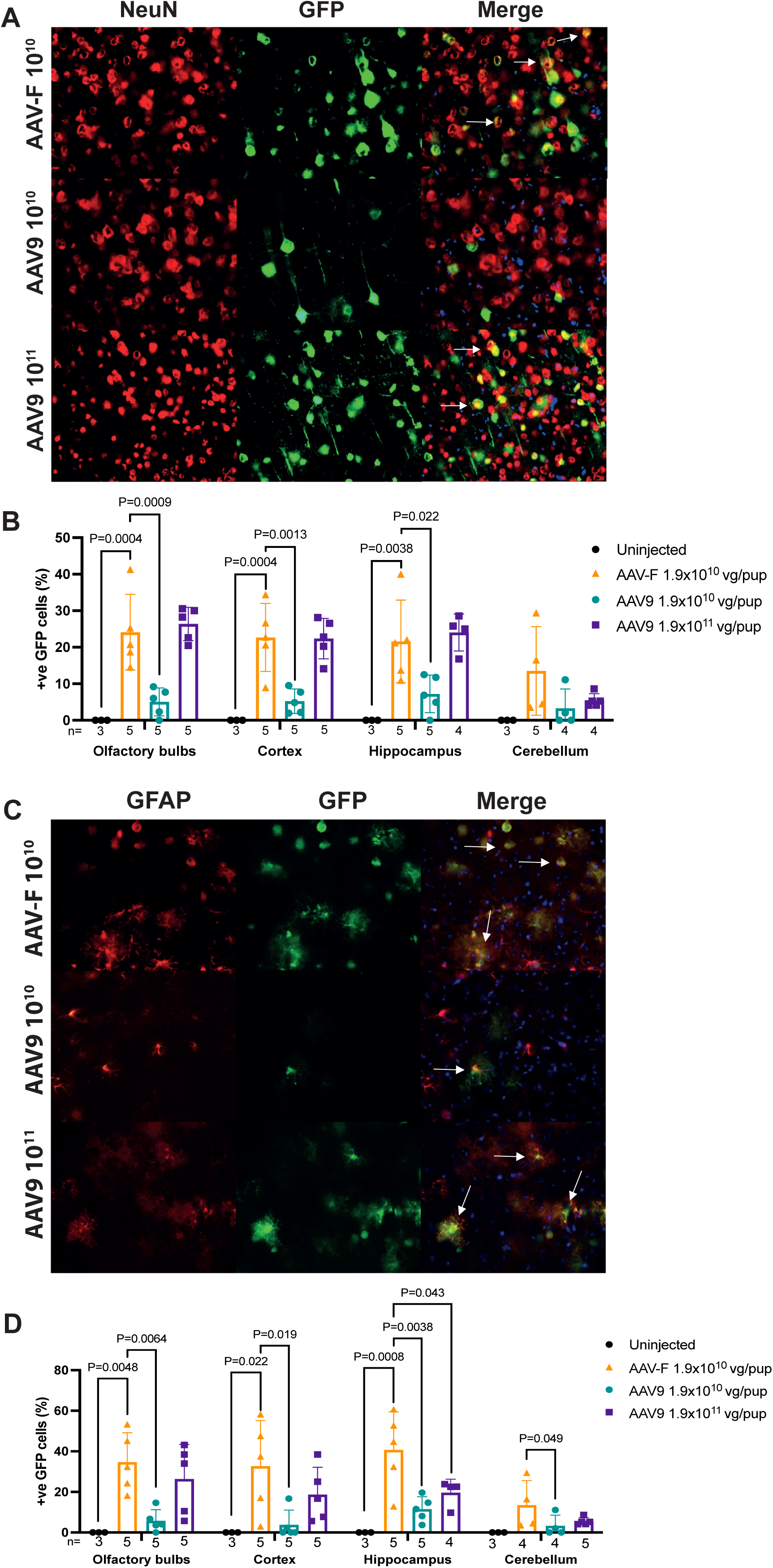
Co-localisation and quantification of neuron and astrocyte targeting of AAV-F. **(A)** Representative images co-localisation of GFP and NeuN. Images were taken at 400X magnification. Scale bar, 100µm. **(B)** Quantification of GFP and NeuN positive cells. One-way ANOVA with Dunnett’s multiple comparisons. **(C)** Representative images co-localisation of GFP and GFAP. Images were taken at 400X magnification. Scale bar = 100µm. **(D)** Quantification of GFP and GFAP positive cells. One-way ANOVA with Dunnett’s multiple comparisons.

**Fig. S7.**
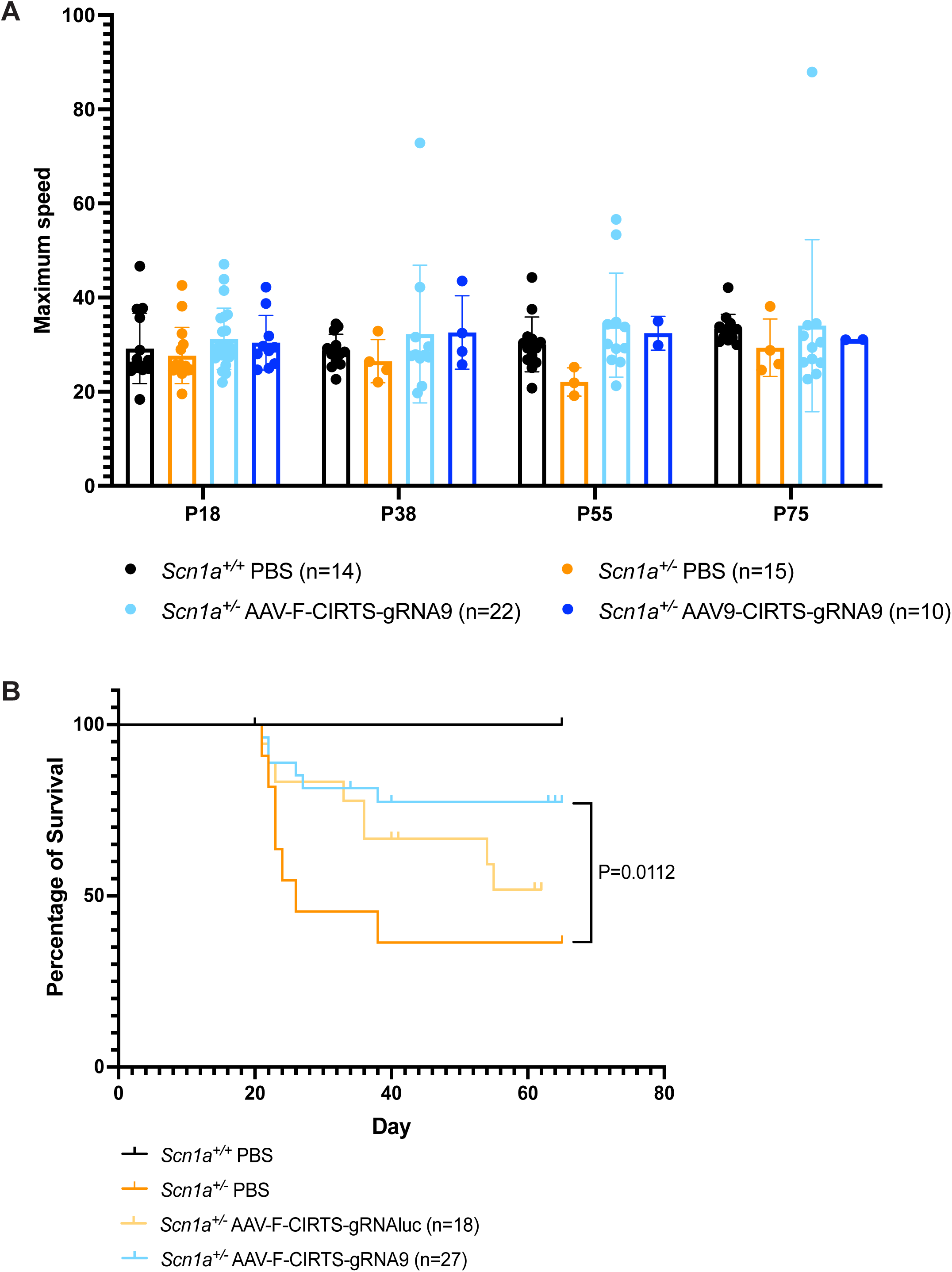
Maximum speed of AAV-F/9-CIRTS-gRNA9 mice and survival curve for AAV-F-CIRTS-gRNA9. (A) Open field maximum speed assessment at 4 different timepoints. Two-way ANOVA with Dunnett’s multiple comparisons. (B) Kaplan-Meier survival curves of treated mice with percentage survival shown. Log-rank (Mantel-Cox) test.

**Fig. S8.**
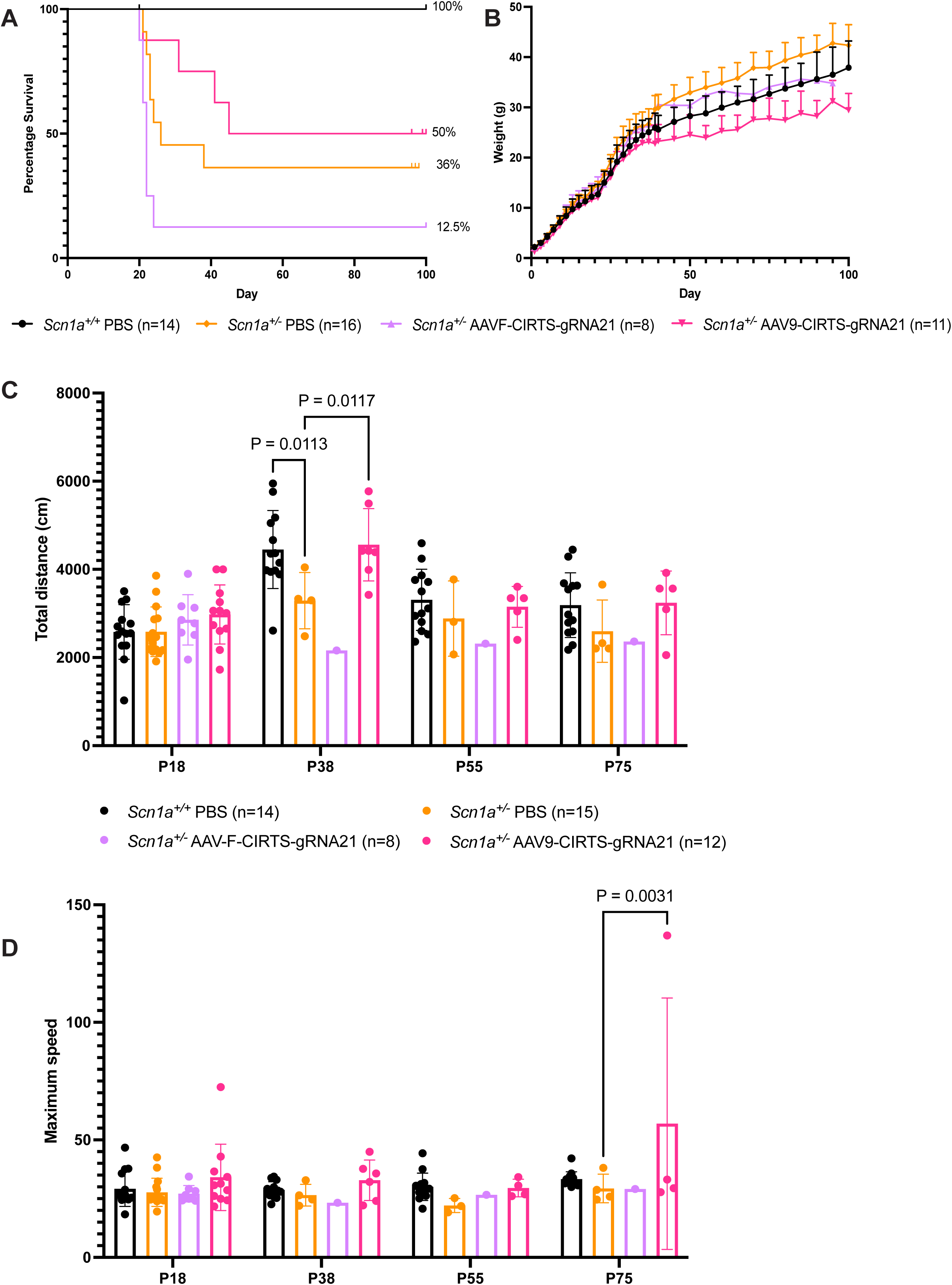
Neonatal AAV-F/9-CIRTS-gRNA21 treatment. *Scn1a^+/-^* mice were administered either AAV-F-CIRTS-gRNA21 at 7×10^10^ vg/pup or AAV-9-CIRTS-gRNA21 at 7×10^11^ vg/pup, with PBS-treated *Scn1a^+/-^*and *Scn1a^+/+^* littermate controls. (A) Kaplan-Meier survival curves of treated mice with percentage survival shown. Log-rank (Mantel-Cox) test. (B) Weights of treated mice. AAV-9-CIRTS-gRNA21 was significantly different from *Scn1a^+/-^* PBS mice at P90 only (P=0.0390), but only female mice were alive in the treatment group at this time (100% female at P100 vs 25% female in *Scn1a^+/-^*PBS). Mixed-effects analysis with Geisser-Greenhouse correction. (C) Total distance and (D) maximum speed read-outs from open field analyses at 4 different timepoints. Two-way ANOVA with Dunnett’s multiple comparisons.

**Fig. S9.**
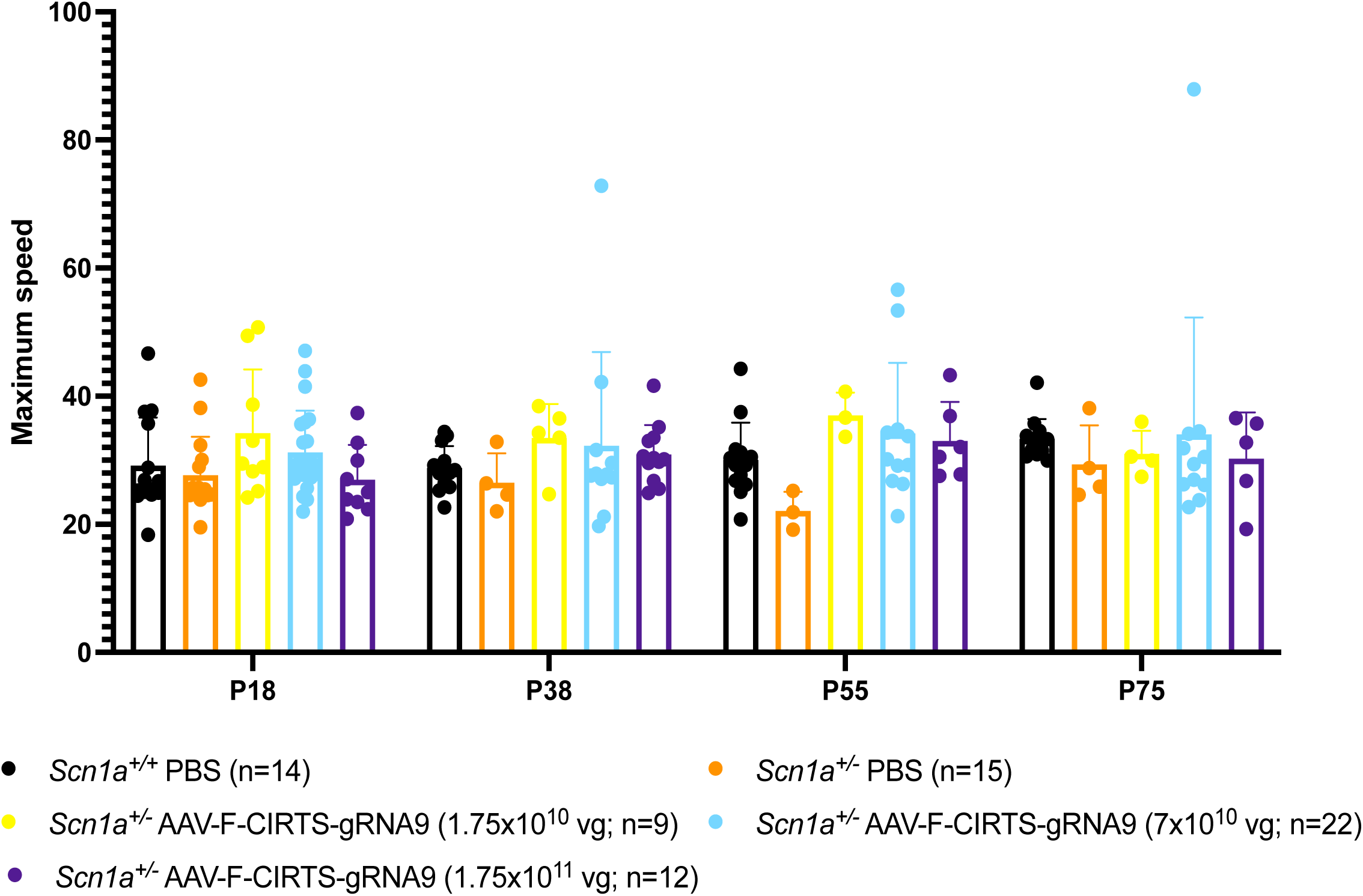
Maximum speed of AAV-F-CIRTS-gRNA9 dosage. 4 different timepoints. Two-way ANOVA with Dunnett’s multiple comparisons.

**Fig. S10.**
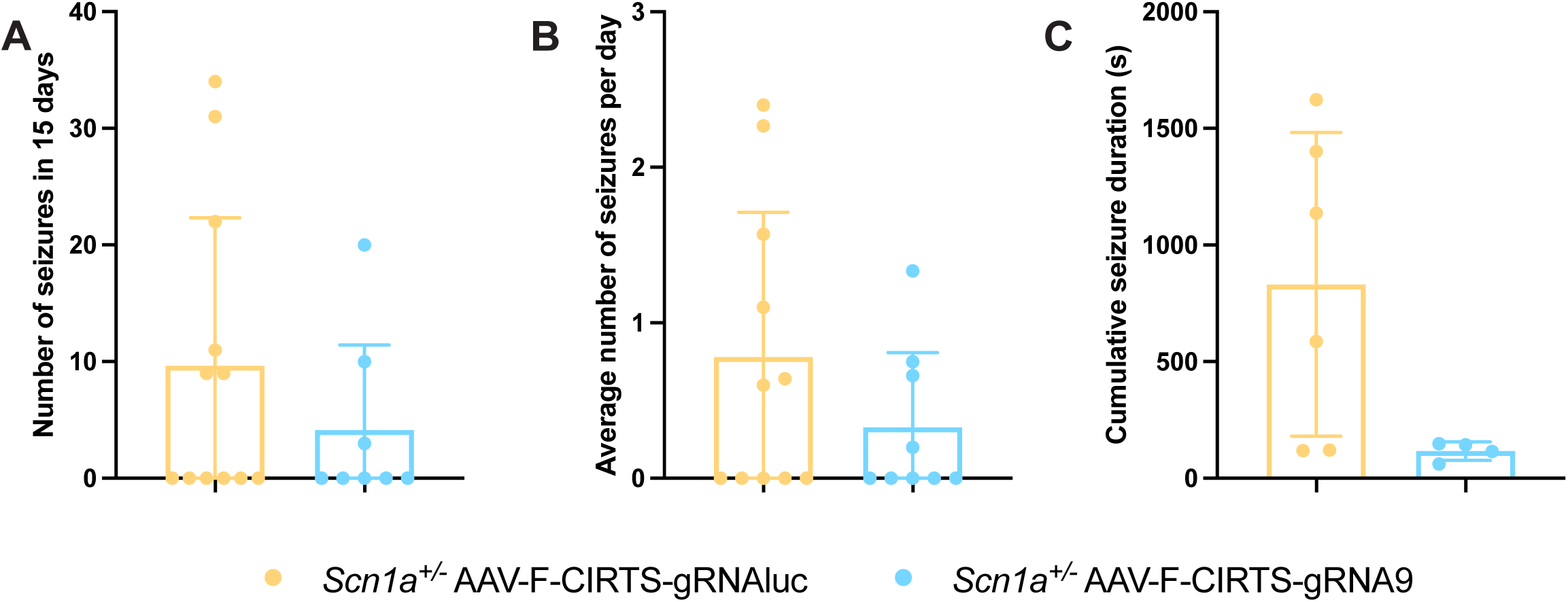
Seizure monitoring in AAV-F-CIRTS-gRNA9 treated DS mice. (A) Total seizures measured in 15 days. Mann-Whitney test. (B) Average number of seizures per day. Unpaired, two tailed t-test (C) Cumulative seizure duration. Unpaired, two tailed t-test.

**Fig. S11.**
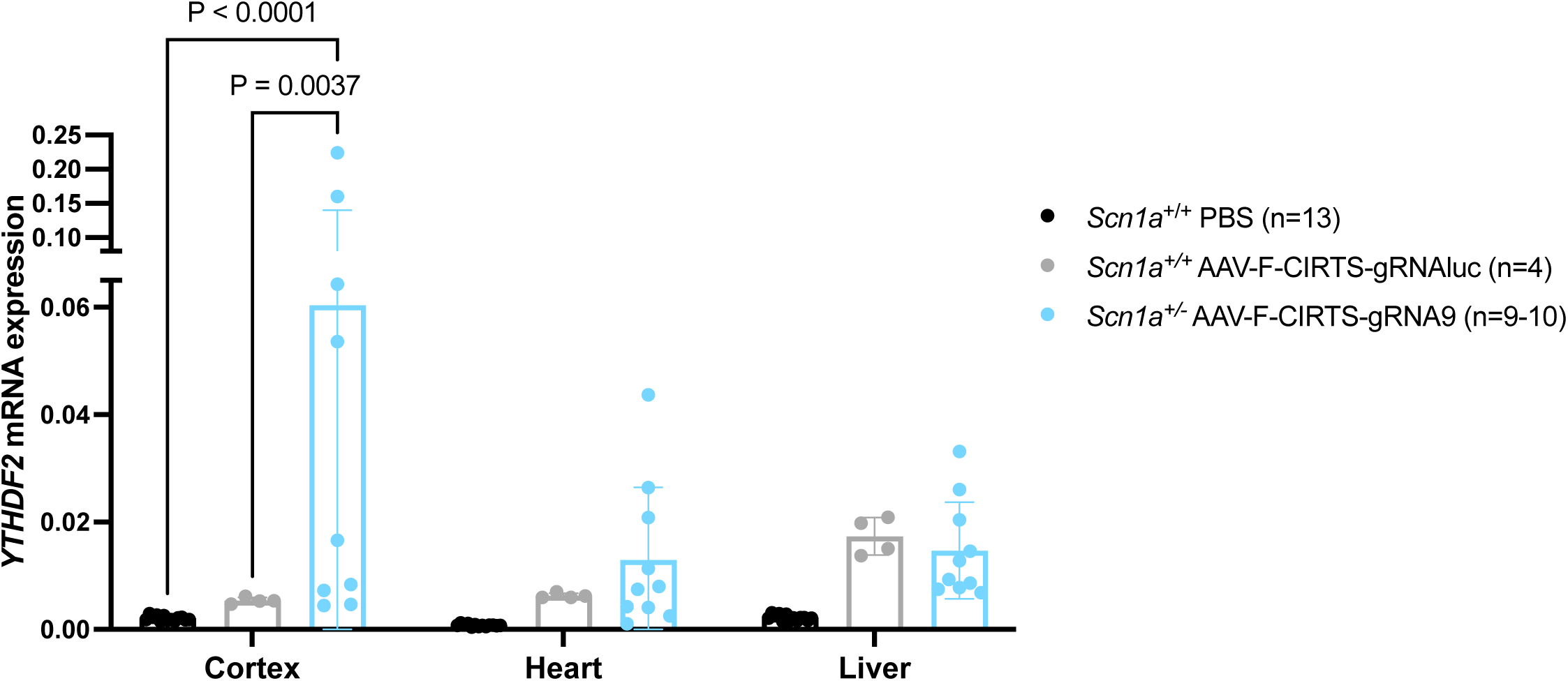
*YTHDF2* gene expression analysis. P100 cortex, heart and liver. Two-way ANOVA with Tukey’s multiple comparisons.

**Fig. S12.**
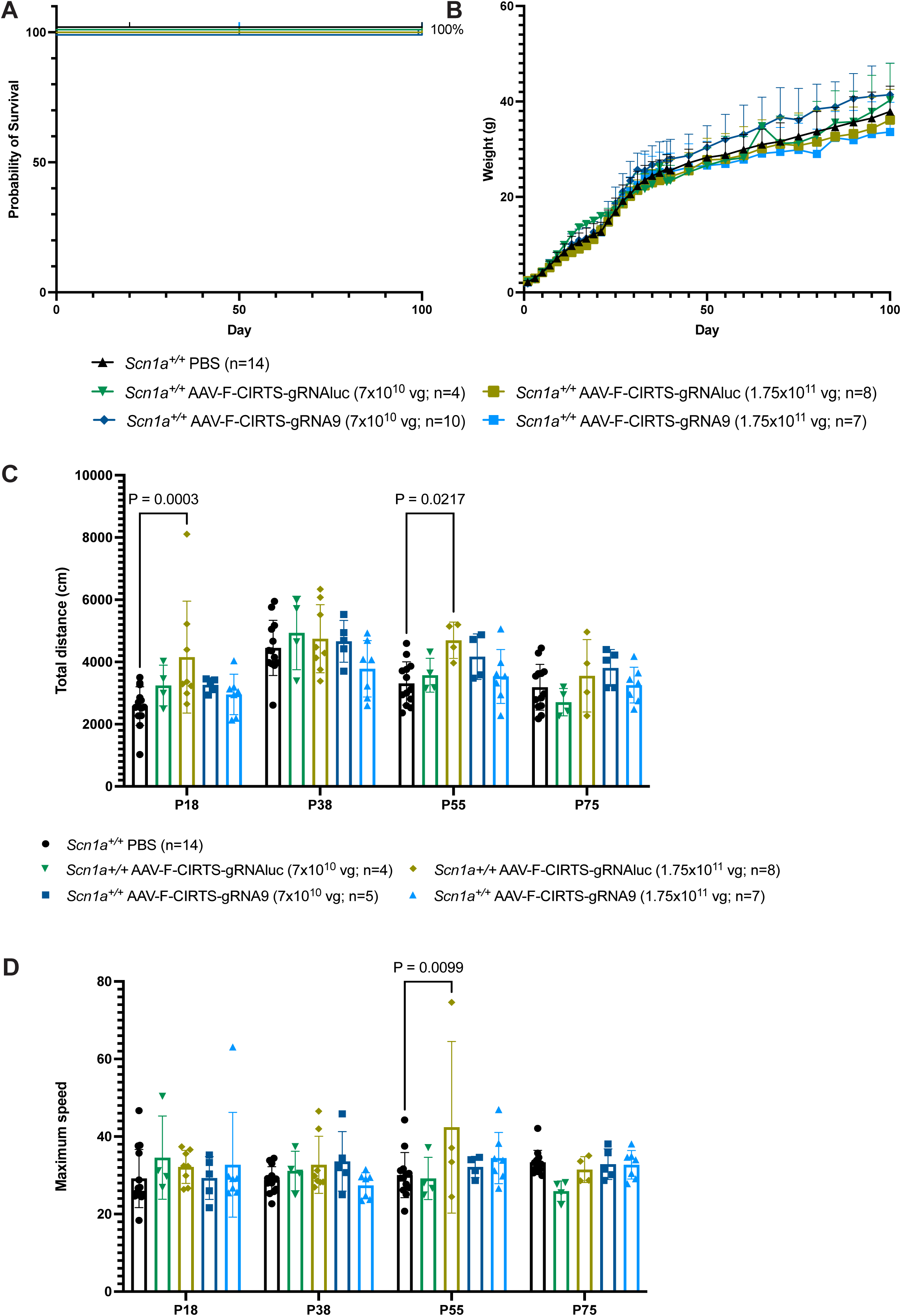
AAV-F-CIRTS-gRNA9/luc administration to healthy mice. *Scn1a^+/+^*mice were administered either AAV-F-CIRTS-gRNA9 or AAV-F-CIRTS-gRNAluc at the optimum therapeutic dose 7×10^10^ vg/pup, or 1 dose higher at 1.75×10^11^ vg/pup to assess signs of toxicity, compared to PBS littermates. (A) Kaplan-Meier survival curves of treated mice with percentage survival shown. Log-rank (Mantel-Cox) test. (B) Weights of treated mice. One-way ANOVA with Dunnett’s multiple comparison. (C) Total distance and (D) maximum speed read-outs from open field analyses at 4 different timepoints. Two-way ANOVA with Dunnett’s multiple comparisons.

